# Quantitative machine learning of protein interactions reveals the multiscale organization and molecular syntax of signaling networks

**DOI:** 10.64898/2026.09.25.754499

**Authors:** Julia R. Rogers, Loren P. Cardani, Neel H. Shah, Mohammed AlQuraishi

## Abstract

Cells employ dense networks of transient protein–protein interactions mediated by modular peptide-binding domains and unstructured peptidic motifs for high-fidelity information processing. How these networks physically execute computations through protein interactions governed by complex intra- and intermolecular mechanisms remains indiscernible from current, sparse and non-quantitative, maps of the human interactome. Here, we introduce a quantitative statistical mechanical modeling (QSM) approach for machine learning domain–peptide affinities with experimental-level accuracy. Leveraging a new, principled algorithm for data harmonization and a biophysically informed neural network architecture, QSM learns to predict dissociation constants directly from amino acid sequences with calibrated confidence. We use QSM to construct the first quantitative drafts of human signaling networks and study these networks across three physical scales—recognition mechanisms of modular binding domains, combinatorial logic of multi-dentate proteins, and pathways inferred from *de novo* inference of protein interaction networks. We find that (i) modular domains, based on their binding preferences, selectivities, and strengths, fall into a limited number of biophysical equivalence groups, (ii) those domains, along with peptidic motifs, are “syntactically” combined within proteins to yield multivalent recognition mechanisms, and (iii) the organization of cellular function can be traced back to algorithmically detectable modules induced by domain-mediated interactions. In aggregate, these analyses instantiate a tractable roadmap towards a comprehensive, mechanistic, and simulatable articulation of the systems biology of signaling.

## Introduction

Cellular physiology and development rely on coordinated, dynamic interactions between thousands of molecularly diverse proteins. Large-scale human interactome atlases, ^1,2^complemented by advances in protein structure prediction, ^3–5^have begun to systematically map protein–protein interaction (PPI) networks and reveal molecular mechanisms underlying cellular organization and behavior in both physiological and pathological states. However, current proteome-scale maps remain highly incomplete and non-quantitative drafts (covering an estimated 2-11% of all pairwise human PPIs ^1^with binary readouts) that cannot explain phenotypic outcomes decided by competitive interactions between proteins with different binding strengths or cellular abundances. ^6,7^

Consistently missing from these maps are numerous weak PPIs mediated by modular peptide-binding domains (PBDs) that recognize peptide sequences, often described as short linear motifs (SLiMs), ^8,9^enriched in intrinsically disordered regions of proteins. Despite SLiMs being short, degenerate sequences present in high-abundance across the proteome (estimated at over 100,000 in humans), ^10,11^PBDs recognize their cognate binding motifs with the fidelity necessary to form effective signaling and regulatory networks. Variations in PBD–peptide binding strength can tune the timing and extent of protein phosphorylation, degradation, localization, and assembly events that collectively determine physiological responses to cell stress, ^12^cell cycle progression, ^13,14^and T cell activation, ^15^to name a few examples. Quantitative differences in PBD binding affinities thus underpin how selectivity and promiscuity are balanced across signaling networks to enable precise control of individual cellular pathways while simultaneously directing pathway crosstalk to orchestrate systems-level behaviors.

Quantitative measurements of PBD–peptide interaction affinities, however, remain scarce. Their weak binding strengths, characterized by dissociation constants in the micromolar range, ^16^challenge experimental detection and necessitate the use of methods with throughput limited to tens or hundreds of interactions, ^17^such as isothermal calorimetry and surface plasmon resonance, or ones requiring specialized equipment, such as holdup ^18,19^and microfluidic platforms ^20^. Experimental efforts to resolve the SLiM-mediated interactome, instead, commonly leverage cell-based surface display ^9,21,22^to screen proteome-scale libraries for interactions. These screens uncover tens of thousands of interactions but fail to quantify absolute thermodynamic constants needed to universally compare binding strengths. ^23^

This paucity of quantitative affinity measurements has stymied computational approaches, partly explaining why machine learning has yet to transform our ability to predict binding affinity from protein sequence as it has our ability to predict structure. ^24–26^The success of machine learning-based structure predictors at modeling well-structured PPIs, moreover, insufficiently transfers to transient PBD-mediated interactions, ^5,27,28^with their characteristic sensitivity to subtle sequence variation and dependence on post-translational modifications (PTMs) ^10^. Tailoring machine learning models to individual PBD families remains a prevailing strategy for capturing these effects, albeit with PTMs represented only implicitly, ^29–31^ even though this limits existing models to a few, well-characterized PBD families (8 out of an estimated 200 in human ^11^) and renders affinity prediction infeasible.

Lacking quantification at proteome-scale, conceptual frameworks for understanding cell signaling have had to resort to one of three archetypical simplifications: (i) binary networks that approximate the scale of regulatory systems ^1,2^but fall short of providing interpretable insights or executable simulations of signaling logic, (ii) narrowly scoped models of well-studied pathways that permit dynamical simulation ^32,33^yet fail to capture the systems behavior crucial to cell function, or (iii) phenomenological models of biological computation that reveal possible mechanisms ^34,35^ but remain agnostic to natural sequence variation. We argue that key to elucidating biological information processing is the development of techniques that broadly and quantitatively predict (or measure) PPIs to yield models that simultaneously capture biological complexity and enable mechanistic reasoning.

A potential approach could exploit the inherent modularity of signaling ^36–38^and build understanding across spatial scales: from sequence-specific motif recognition by individual binding domains, to multivalent protein binding governed by combinatorial interactions—what can be understood as molecular syntax encoding rules for protein regulation and modulation—to network topology and large-scale organization, and ultimately to the logic computed by these networks. Here, we take a step towards this aim with a quantitative statistical mechanical modeling (QSM) approach that yields a series of telescoping machine learned physical models. At the base level, QSM quantitatively predicts PBD–peptide interaction affinity directly from protein sequence. It introduces a principled framework for integrating heterogeneous experimental binding measurements that overcomes data scarcities to model biochemically and structurally diverse PBD families and the consequences of phosphorylation, methylation, and acetylation on binding. Through both retrospective and prospective evaluations, we demonstrate that QSM predicts the dissociation constants of PBD–peptide interactions with errors comparable to experimental methods, and that its self-estimates of prediction accuracy indicate when these values can be trusted. Prediction of absolute thermodynamic constants enables QSM to synthesize domain-level affinities into physical models of modular proteins comprising multiple PBDs and/or SLiMs, yielding a platform for multivalent protein interaction discovery that substantially outperforms existing computational and experimental approaches and that can uncover biophysical interaction mechanisms, the molecular syntax of signaling, *en masse*.

Applied proteome-wide, QSM reveals principles and organization underlying PBD-mediated signaling across multiple scales: at the domain level, (i) we distill the functional consequences of PBD sequence variation into a classification system of biophysical equivalence groups exhibiting similar binding preferences and (ii) identify strength and selectivity as the primary, and largely orthogonal, axes of variation within PBD families; at the network level, (iii) we construct—starting from protein sequence information alone—to our knowledge the first quantitative human interactome with broad coverage of PBD-mediated PPIs, and (iv) characterize molecular interaction mechanisms that organize PBD-mediated PPIs into biological pathways. While short of producing immediately executable networks, QSM nonetheless lays a foundation for a bottom-up and mechanistic approach to studying cell signaling by fusing the complementary advantages of machine learning and physics-informed modeling.

## Results

### Learning from heterogeneous binding data

#### Model overview

QSM integrates information from heterogeneous binding assays and across PBD families using novel learning objectives and network architectures to overcome data disparities. These innovations stem, in no small part, from formulating PBD–peptide affinity prediction statistical mechanically. Statistical mechanics connects macroscopic, thermodynamic properties to probability distributions over microscopic states, relating dissociation constants (*K*_D_’s) to exponential changes in free energies upon binding via ratios of partition functions. By machine learning an empirical approximation to the free energy function of binding—instead of directly predicting affinities themselves— QSM naturally integrates experimental readouts from assays that have traditionally been modeled as distinct within a principled learning paradigm, which we term Single Objective, Multiple Data (SOMD) learning. We design the QSM neural network free energy function to learn a mapping from the amino acid sequences of a PBD and peptide to binding free energy that satisfies established biophysical constraints, introducing a network primitive that facilitates learning of physiological affinity values. These strategies are combined with self-estimates of accuracy, informed model ensembling and data sampling across PBD families, protein language model (pLM) sequence representations, and explicit modeling of post-translational peptide modifications (Fig. 1) to enable transfer of knowledge from PBD families with abundant binding data to ones with limited data—or even lacking training data entirely—in a trustworthy manner.

**Fig. 1.**
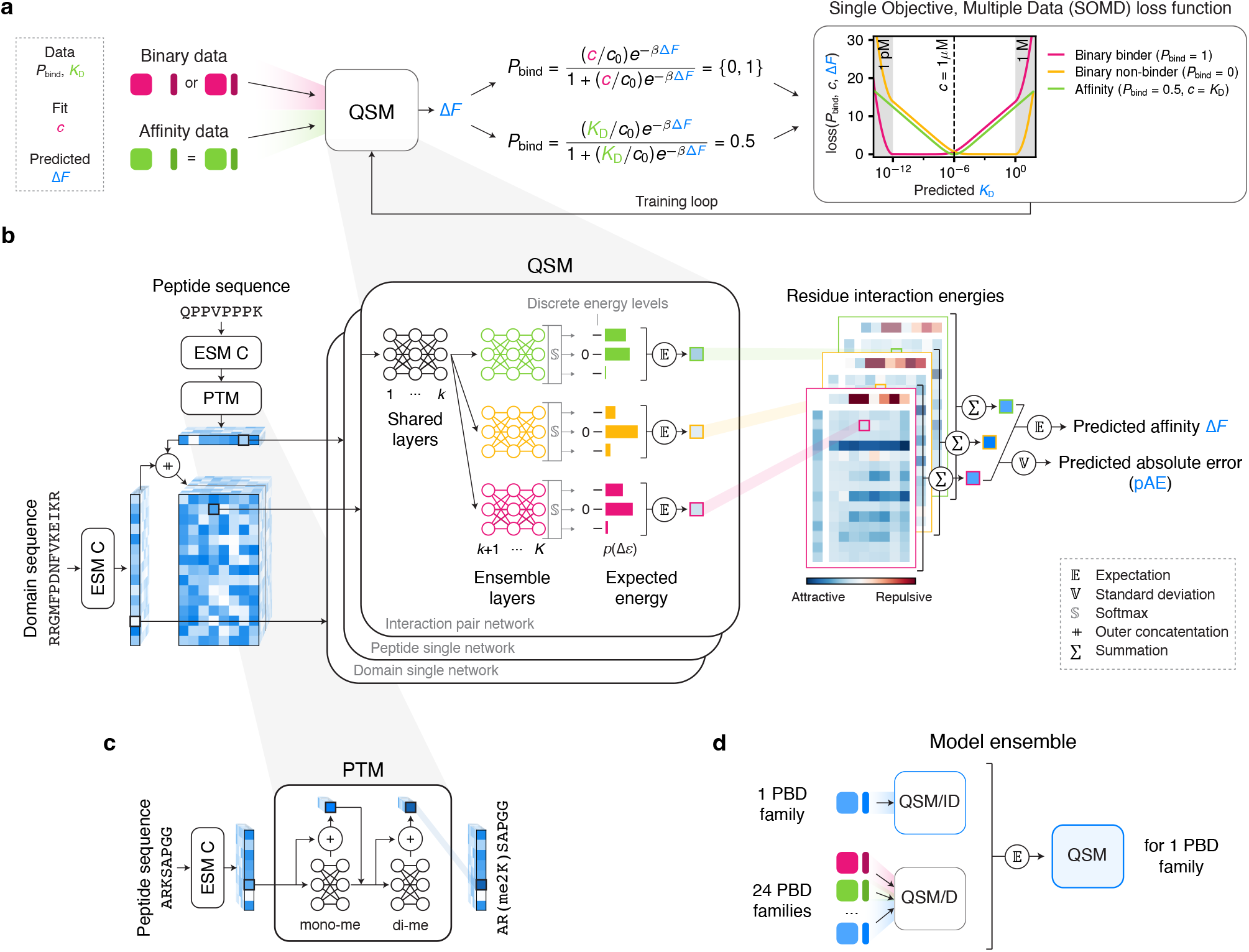
QSM training algorithm and architecture facilitate learning from heterogeneous binding data. **a.** QSM simultaneously learns from binary binding measurements and quantitative a!nities using Single Objective, Multiple Data (SOMD) learning. **b.** QSM predicts the change in free energy upon binding from input domain and peptide sequences. Sequences are first embedded using ESM C and then processed by three shallow neural network ensembles to estimate residue-level contributions to the binding energy as expectations over discrete energy values, finally outputting a predicted a!nity and predicted absolute error of this prediction. **c.** Representations of post-translationally modified peptide residues are updated using a series of residual networks for each modification modeled by QSM (phosphorylation, methylation, and acetylation). **d.** QSM ensembles a model trained on one PBD family (QSM/ID) with a model trained on all PBD families (QSM/D).

#### Single objective, multiple data learning

QSM models are trained on both copious, qualitative experimental data readily acquired through high-throughput assays, such as peptide arrays and phage-display, and scarce, quantitative affinity measurements mostly obtained through low-throughput assays, such as isothermal titration calorimetry and fluorescence polarization. Traditionally, predicting the read-outs of qualitative binding assays, which only indicate if an interaction is detected (‘binder’) or not (‘non-binder’), is formulated as a binary classification problem, whereas quantifying affinities is considered a regression problem. However, this is a false dichotomy that limits the amount of data available to train quantitative affinity predictors. Both types of experimental readouts, in fact, provide empirical estimates of macroscopic binding probabilities (*P*_bind_): Conclusive ‘binder’ and ‘non-binder’ calls report near certain observation of the bound (*P*_bind_ = 1) and unbound state (*P*_bind_ = 0), respectively, when assayed at a given ligand concentration; measured *K*_D_’s report the ligand concentration at which both states are equally likely 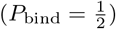. With specification of ligand concentration, macroscopic binding probability is then determined by the change in free energy upon binding. Thus, by predicting this free energy difference, QSM outputs can be directly transformed into both types of experimental readouts. This enables optimization of all model parameters to maximize the likelihood of these two experimental data sources simultaneously, with model agreement to each evaluated using the same loss function (Fig. 1a). The only difference is that the presumed ligand concentration is equated to the *K*_D_ when training on affinity data but must be fit when training on binary measurements.

By training QSM models end-to-end on multiple, distinct data streams using equivalent objectives, SOMD learning avoids the need for assay-specific transformations or trainable network layers to facilitate knowledge transfer across data domains. SOMD learning shares a core motivating idea with multitask learning, ^39^ namely that training models to concurrently solve multiple tasks can improve generalization and data efficiency. However, the two paradigms differ in implementation. Common multitask learning setups require separate task-specific prediction heads on top of shared representational trunks and often combine incommensurate losses. Since each prediction head can only be trained with its own task specific dataset, this may bottleneck learning when there are severe data imbalances across tasks, such as in our case (binary PBD–peptide binding measurements outnumber *K*_D_’s 100-fold). SOMD avoids this. We, nevertheless, still include a hyperparameter to controllably trade-off learning from small-scale, higher-quality affinities versus large-scale, noisier binary measurements.

#### Neural binding free energy function

Our second innovation increases data efficiency by formulating the functional form of the QSM neural free energy function based on known biophysical principles (Fig. 1b). Firstly, epistasis observed in experimental sequence-affinity landscapes of multiple protein–peptide interactions is largely explained by additive effects of single-residue mutations. ^40,41^This motivates our decomposition of the change in energy upon binding into additive contributions from individual residues and interactions between pairs of residues, one in the PBD and one in the peptide. Nevertheless, higher-order interactions must be accounted for to fully explain observed sequence-affinity relationships, ^41,42^and we describe our approach to model these effects in a computationally tractable way next. Secondly, only a few key residues, commonly referred to as hotspots, typically contribute most of the binding energy, with single mutations altering binding free energies by at most a few kcal/mol. ^40,41^This motivates our design of a network primitive to constrain the magnitudes of each energetic term and to encourage sparsity within residue-level energy maps. In QSM, neural networks dedicated to each energetic term compute binding energies as expectations over bounded, discrete energy differences centered around zero kcal/mol, incorporating this prior molecular-level knowledge directly into the architecture. While these two architectural design choices serve to regularize the model, they also yield an interpretable model whose predictions can be explained in terms of residue-specific contributions to binding. Thirdly, PBDs and peptides evolved to interact weakly, with affinities in the micromolar range. ^10^We precondition the model to predict *K*_D_’s within the range measured experimentally prior to training, and include an additional penalty term in our objective function to softly constrain predictions to physiological values when training on binary measurements (Fig. 1a).

#### Post-translationally modified pLM representations

Our third contribution enables information transfer across PBD–peptide interactions by leveraging the ESM C pLM ^43^ to map amino acid sequences to a universal representation space that organizes sequences according to physicochemical, structural, and functional similarities. Parameterizing the QSM free energy function by residues’ ESM C representations (Fig. 1b) allows the model to reason about molecular biophysics shared both within and across PBD families. Moreover, because each residue’s representation depends on all other amino acids in the sequence, QSM learns to contextualize single residue differences in terms of the rest of the protein sequence, implicitly modeling higher-order interactions that shape sequence-affinity landscapes ^41,42^despite explicitly computing only first-order interaction energies. As such, a single mutation can elicit changes in QSM residue-level energy maps at multiple positions anywhere in the sequence.

ESM C (and almost all existing pLMs ^44^) only represents sequences of the canonical 20 amino acids; however, many PBDs recognize PTMs. To explicitly model the effects of PTMs on binding, QSM updates the ESM C representations of each modified peptide residue using a cascading series of residual networks (Fig. 1c). Each of these networks learns to represent the addition of a single chemical group (*e.g.*, phosphate) to any amino acid, and is trained to iteratively apply updates to encode higher-order modification states (*e.g.*, di- and tri-methylation). This chemically informed design pools data for rarely measured modifications with common ones to train each network.

#### Confidence estimation

Our fourth contribution does not directly improve model generalization but addresses issues encountered when models are applied to regimes poorly covered by or entirely outside their training data distribution, which is bound to occur for QSM models constructed for data-limited PBD families. In these regimes, machine learning models are susceptible to making overly confident, wrong predictions. To mitigate this pathology, QSM quantifies its own uncertainty to indicate which predicted affinities can be trusted. Similar confidence estimates provided by AlphaFold, ^24^ for instance, have enabled researchers to propose valid hypotheses for a protein’s function based on its predicted structure even for protein classes underrepresented in its training set. ^45^ In another example, confidence estimates allowed researchers to effectively prioritize small molecule kinase inhibitors from a library of over a thousand compounds for experimental testing, achieving a 90% hit rate with a model trained on less than 100 compounds. ^46^In later sections, we likewise leverage QSM confidence estimates for model-guided hypothesis formulation and PBD–peptide interaction discovery.

QSM associates each of its affinity predictions with a confidence estimate that quantifies its own predicted absolute error (pAE). This is achieved by training an ensemble of neural network free energy functions and computing the pAE as the standard deviation of the individual predicted affinities. QSM specifically uses shallow ensembles, where the last layers of the network are specific to each ensemble member and early layers are shared among them (Fig. 1b), for their implementation simplicity and computational efficiency. ^47^When training on affinities, we include an additional mean-squared-error loss on the pAEs to improve the quantitative calibration of QSM confidence estimates.

#### Model ensemble

Our fifth contribution facilitates knowledge integration across PBD families with disparate dataset sizes by employing model ensembling to address failure modes stereotypical of either data-poor or data-abundant PBD families. Indeed, ensembles succeed at improving performance, generalization, and robustness when their component models make different errors, ^48^and we observe this when QSM models are trained in two ways. First, we train independent models for each of the 24 PBD families that have a mix of affinity, binary positive, and binary negative measurements in a dataset of over 3 million total PBD–peptide binding measurements, whose curation and content we detail in a concurrent manuscript. ^49^We refer to these as QSM for independent domains (QSM/ID). Second we train a single model on data pooled across these 24 PBD families, sampling measurements from each family according to a power-law distribution to best balance learning across families with orders-of-magnitude different dataset sizes. We refer to this model as QSM for domains (QSM/D). QSM/ID performs well on PBD families with abundant data (over 10^5^training measurements), whereas QSM/D performs better for families with limited data (less than 10^3^training measurements) that benefit from data pooling to increase the overall size of the training set (Fig. S1). We then ensemble each QSM/ID model with QSM/D to construct QSM models for each of the 24 PBD families (Fig. 1d). QSM, thus, combines general knowledge of PBD–peptide interactions captured by QSM/D with family specific information learned by QSM/ID, outperforming both QSM/ID and QSM/D for 17 (71%) of these PBD families.

Because QSM models are, nevertheless, still specific for only one of 24 PBD families, we devised an uncertainty-guided ensembling approach to predict interactions involving PBD families outside this set. Specifically, among the pool of available pretrained QSM/ID models, we ensemble the lowest pAE model with QSM/D to yield QSM predictions for these out-of-distribution interactions.

### QSM quantitatively predicts domain binding a!nity from sequence

We first evaluated QSM on each of the 24 PBD families in its training distribution using 5-fold cross-validation, in which unique PBD–peptide pairs were randomly assigned to each fold. QSM predicts *K*_D_’s within an order of magnitude of experimental values for 95% of interactions, on average achieving 1 kcal/mol accuracy, and quantitatively correlates with experimental measurements, obtaining Pearson correlation coefficients (*R*) above 0.7 for half of the 24 PBD families (Fig. 2a). We also evaluated QSM on classifying PBD–peptide pairs as binders or non-binders to assess it on a larger, more sequence-diverse set of interactions. QSM classifies interactions with a median per-family area under the receiver operating characteristic curve (AUROC) of 0.96, and demonstrates strong recall at low false positive rates (exceeding 75% at a 10% false positive rate for nearly all PBD families) (Fig. S2), indicating that QSM can reliably discover PBD–peptide interactions.

**Fig. 2.**
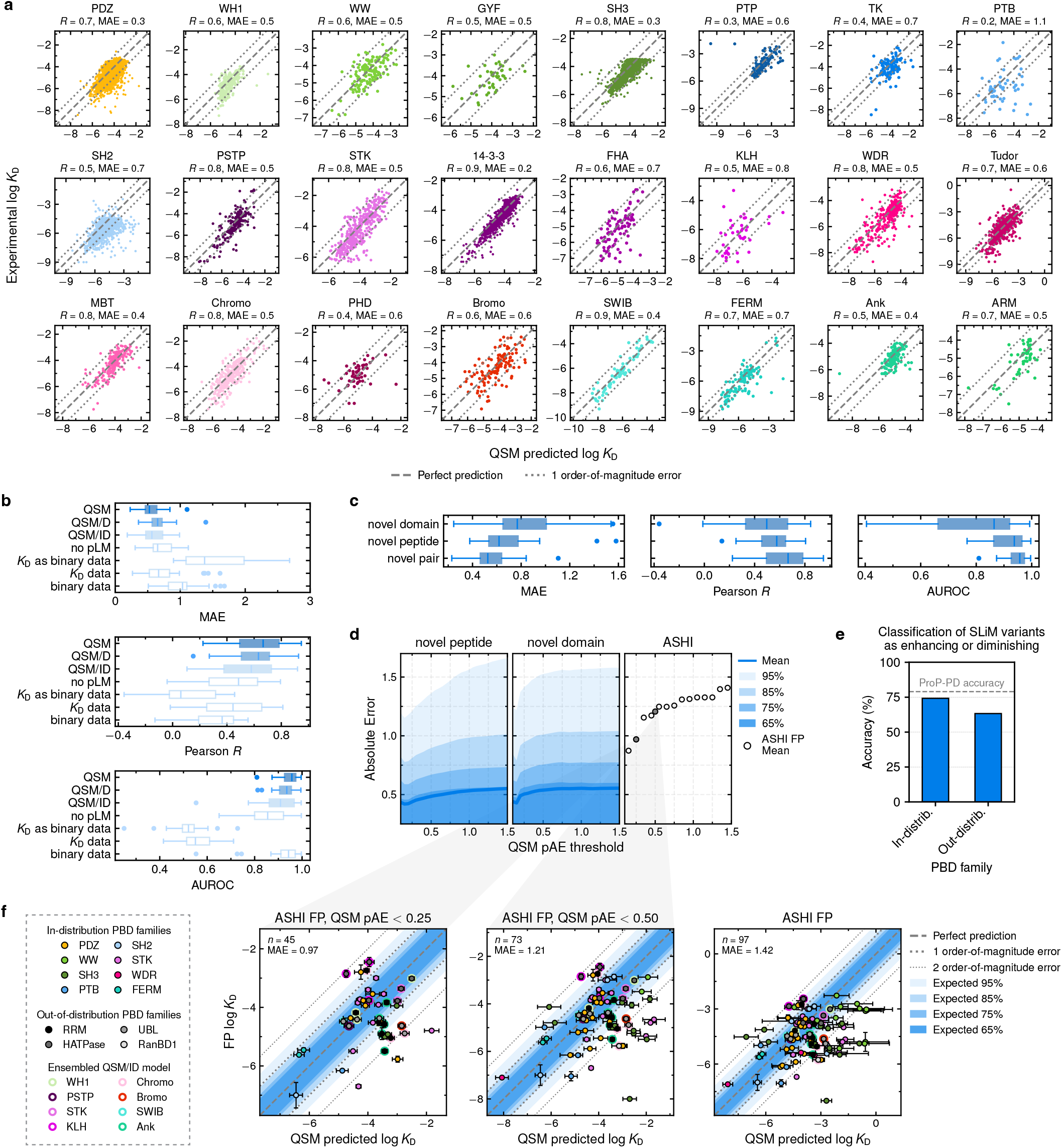
QSM quantitatively predicts a!nity from amino acid sequence. **a.** Experimental vs predicted a!nities for 24 PBD families evaluated in 5-fold cross-validation. **b.** Ablations of QSM components: ESM C is replaced in QSM/ID with learnable amino acid embeddings and rotary position embeddings (“no pLM”), QSM/ID is trained only on interactions with available *K*_D_’s but with binary training labels (“*K*_D_ as binary data”), only on *K*_D_ data, or only on binary data. Box-and-whisker plots of per-family mean absolute error (MAE) and Pearson correlation coe!cient *R* of log *K*_D_ predictions, and area under the receiver operating characteristic curve (AUROC) for classifying binders from non-binders. MAE clipped to 3 with “no pLM” MAE *>* 3 for one family, and “*K*_D_ as binary” MAE *>* 3 for three families. **c.** Box-and-whisker plots of per-family QSM performance evaluated in 5-fold cross-validation on held-out test domains (“novel domain”), peptides (“novel peptide”), or interactions (“novel pair”). **d.** True absolute error of QSM predicted log *K*_D_ versus QSM predicted absolute error (pAE) for data splits in **c** and a!nities reported in ASHI ^9^. **e.** Accuracy of QSM at classifying human short linear motif (SLiM) mutations as enhancing (reduced *K*_D_ relative to wild type) and diminishing (increased *K*_D_) of binding to PBDs from both in- and out-of-distribution families, with proteomic peptide phage display (ProP-PD) providing ground truth. Only predicted a!nity changes exceeding the prediction error are classified. **f.** *K*_D_’s determined using fluorescence polarization (FP) competition assays in ASHI versus QSM predictions at di”erent pAE thresholds.

QSM pAE values indicate when a predicted affinity can be trusted (Fig. S3 and 2d). While QSM pAE, itself, typically underestimates the true absolute error, affinities with low pAE are more trustworthy: 87% of the predicted affinities with pAE *<* 0.25, which we consider high confidence, agree with experimental *K*_D_’s within a factor of 4, whereas the same proportion of predictions with greater pAE agree within only a factor of 16.

Ablations of individual QSM components demonstrate that multiple strategies to integrate biophysical knowledge present across heterogeneous binding measurements and diverse PBD families, indeed, contribute to QSM’s accuracy (Fig. 2b and Table S1). Neither data type alone—at the scale currently available—is sufficient to train quantitative sequence-based predictors of PBD binding affinity. Training on only one source reduces median per-family *R* by 24-38% and increases mean absolute error (MAE) by 20-86%. QSM’s quantitative accuracy stems from training on *K*_D_ measurements but benefits from training on larger binary datasets that cover a broader swath of sequence space. Notably, a model trained on *K*_D_ measurements alone, without model ensembling, achieves a median per-family *R* of 0.44, while the QSM model trained on all data achieves a median per-family *R* of 0.67, demonstrating that SOMD leverages binary data to materially improve the quantitative accuracy of the model. ESM C representations, capturing physicochemical, structural, and evolutionary information observed across billions of protein sequences, ^43^ further facilitate reasoning across sequence space. Replacing them with ones learned through supervised training results in performance drops comparable to training on only one data source.

### QSM generalizes to new domains and peptides

We next assessed how well QSM predicts the impact of sequence variation on binding affinity and how well it models PBDs and PBD families outside its training distribution. We emphasize the distinction between PBDs and PBD families, as some of our evaluations test generalization to previously unseen PBDs within a family (”in-distribution”; *e.g.*, an individual PDZ domain) while other evaluations test generalization to unseen PBD families (”out-of-distribution”; *e.g.*, the entire RRM family).

Our first evaluation tested QSM on PBDs or peptides excluded entirely from training (*i.e.*, not assayed against any peptide or PBD in the training set, respectively) to characterize its sensitivity to sequence variation within a PBD family or peptide motif. We note that sequence variation in this evaluation set results from both natural PBD evolution and targeted mutagenesis or positional scanning assays designed to probe biochemical recognition mechanisms. Although performance degrades on held-out sequences (Fig. 2c), QSM predicts *K*_D_’s within an order of magnitude of experimental values for 85% and 83% of interactions involving held-out peptides or PBDs, respectively. QSM generalizes better to new peptides than PBDs, potentially because training datasets more densely cover the space of possible peptide sequences (outnumbering unique PBD sequences by 10-10,000 fold, which can be as few as 4 for a single PBD family). QSM pAE remains informative of high confidence predictions (Fig. 2d), indicating that it can support human decision-making even in novel settings.

Our second evaluation tested QSM on predicting the impact of disease-associated mutations found in human intrinsically disordered regions on PBD–peptide binding. We compared QSM-predicted mutational effects to results from high-throughput proteomic peptide phage display (ProP-PD) that classified mutants as diminishing or enhancing binding to a library of PBDs ^50^spanning 12 in-distribution families and 32 out-of-distribution ones. Results from this experimental study were not compiled in QSM’s training dataset, nor are any assayed mutant-wild type interaction pairs measured in other experimental studies included in QSM’s training set, making this an independent evaluation of generalization. Following a classification protocol that accounts for model uncertainty, which would foreseeably be employed by researchers to predict functional consequences of newly identified sequence variants, QSM predicts variant effects with accuracies approaching that of the ProP-PD screen itself (Fig. 2e). Thus, QSM qualitatively models PBD–peptide interactions with single mutation sensitivity comparable to high-throughput experiments.

Our third evaluation tested QSM on quantifying 97 PBD– peptide *K*_D_’s reported in the Atlas of SLiM-mediated Human protein–protein Interactions (ASHI) ^9^after we completed QSM training. This independent test set includes interactions with *K*_D_’s spanning the nanomolar to millimolar range and mediated by 15 PBDs present in QSM’s training set, 8 PBDs from in-distribution families but absent from the training set, and 5 PBDs from out-of-distribution families, allowing us to assess generalization at three levels of difficulty. Consistent with earlier results, predicted affinities with low QSM pAE are more reliable (Fig. 2d,f). High-confidence predictions agree within an order of magnitude of experimental *K*_D_ values for 72% of interactions mediated by in-distribution PBD families (73% mediated by domains represented in the QSM training set), and 55% mediated by out-of-distribution families, with MAEs mirroring this trend and corresponding to average binding free energy discrepancies of roughly 1.3 kcal/mol (Fig. 2f). Overall, this illustrates that QSM can quantitatively generalize to novel interactions, including those involving out-of-distribution PBD families, although these predictions should generally be regarded with more caution than those involving one of QSM’s 24 training families. Moreover, this evaluation demonstrates that QSM predicts affinities with accuracies rivaling experimental methods—including high-throughput holdup approaches ^51^developed specifically for PBD-mediated interactions—which commonly differ across labs and techniques by 0.6-1 kcal/mol ^52^and can report *K*_D_’s that differ by an order of magnitude, as we observe in our own experimental validations in a later section.

### PBDs form a small number of biophysical groups

We leveraged QSM’s ability to rapidly quantify affinities with accuracies comparable to experiments to investigate how the binding preferences of 2,754 PBDs from the 24 in-distribution families vary across the human proteome. By comparison, high-throughput assays suitable for characterizing the largest PBD families, such as PDZ (275 domains) and SH3 (325 domains), identify key sequence features of domain recognition—and codify them into consensus binding motifs ^8,53,54^—but do not reliably quantify differences in binding affinity that determine promiscuity, selectivity, and specificity ^9^.

Among half a billion predicted affinities for over a million peptidic sites found across the human proteome with physicochemistries similar to known binding motifs, including post-translationally modified sites, we limited our analysis to domains with confidently predicted affinities for a shared set of peptides, each predicted to bind at least two domains, to reliably compare their binding preferences (Fig 3a). We clustered domains within a PBD family based on the similarity of their QSM predicted affinities for each peptidic site to assign them to biophysical groups: Domains that belong to the same group have affinity differences that are nearly indistinguishable from thermal energy fluctuations and, thus, can be considered biophysically equivalent in terms of their binding preferences, while domains that belong to different groups typically have over one order-of-magnitude different *K*_D_’s for the same peptidic site (Figs. 3a, S5, and S6). Across 21 PBD families, we identify 132 biophysical groups, 15-fold fewer unique groups than unique domains (Fig. 3b), indicating that the binding preferences of thousands of domains can be understood using a simple classification system.

**Fig. 3.**
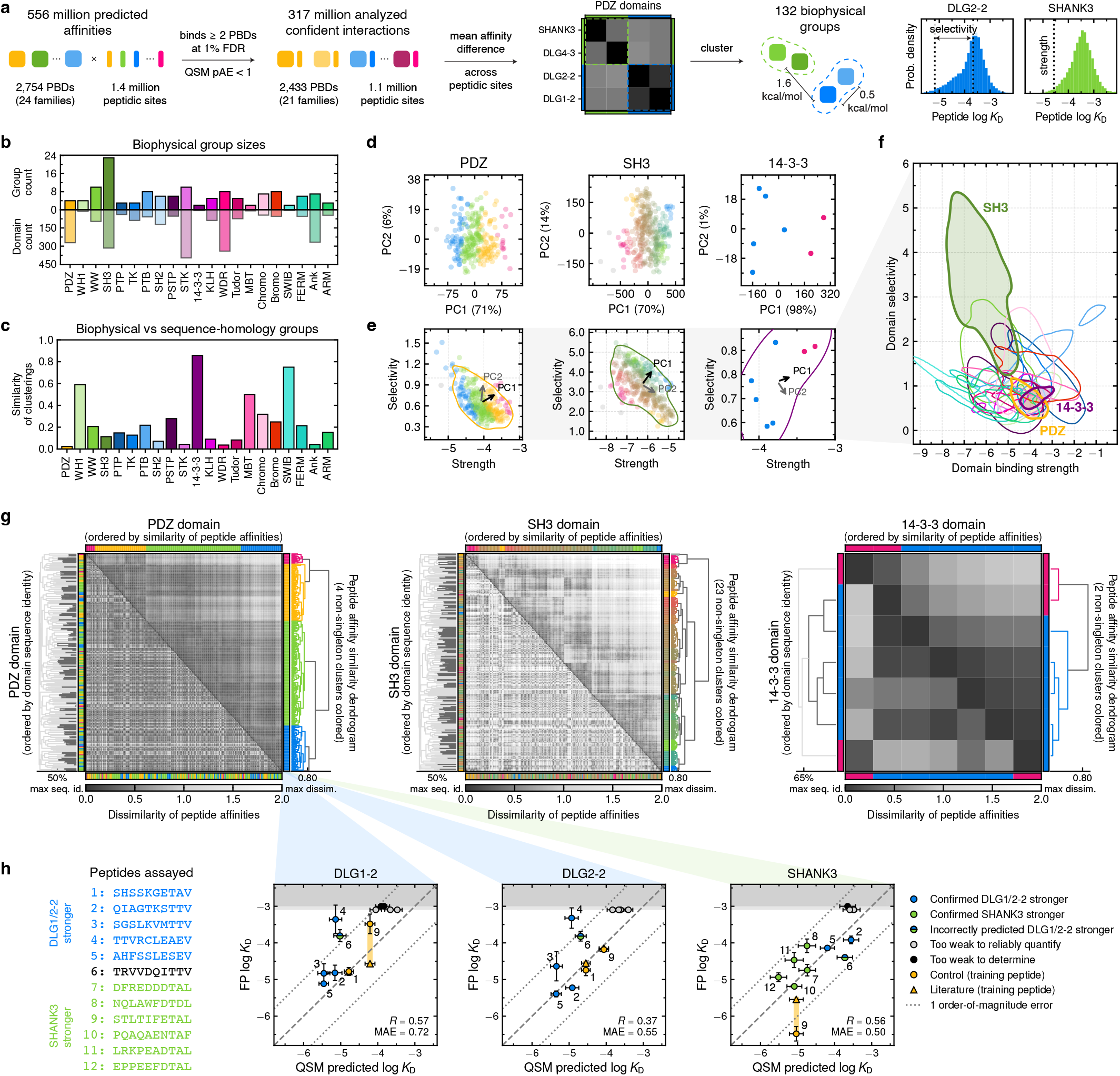
Proteome-wide classification of PBDs into biophysical equivalence groups. **a.** Pipeline to identify biophysical groups. Example a!nity distributions for two PDZ domains classified into groups B (blue) and G (green) illustrate definition of binding strength (top 2% of peptidic sites) and selectivity (di”erence in a!nities between the top 2% and median of peptidic sites). **b.** Number of biophysical groups and PBDs per family. **c.** Maximum concordance of biophysical and sequence-homology groups quantified by an element-centric comparison framework. ^55^ **d.** PCA of PBD–peptide a!nities. PBDs projected onto first 2 PCs (percent variance given in parentheses) and colored by biophysical group. **e, f.** Binding strengths and selectivities of PBDs. Contours outline 90% of the probability mass per family. Arrows in **e** indicate directions of first two PCs determined through least squares regression. **g.** Similarity matrices of peptide a!nities. Upper half is ordered by domains’ similarity of peptide a!nities, and lower half is ordered by their sequence similarities. Colors indicate biophysical group assignments and are assigned based on cophenetic distance. Clustering thresholds indicated on each dendrogram. Domains that cluster together based on sequence identity colored dark gray on dendrogram. **h.** Experimentally determined *K*_D_’s from fluorescence polarization (FP) assays versus QSM predictions for PDZ domains assigned to biophysical groups B (blue) or G (green) against a panel of 12 human proteome-derived peptides. Pearson correlation coe!cient (*R*) and mean absolute error (MAE) computed for interactions with experimentally quantified *K*_D_’s. A!nities that were too weak to reliably quantify are plotted at *K*_D_ = 800*µ*M, the highest concentration tested (light gray region); weaker interactions that could not be reliably detected are plotted at *K*_D_ = 1mM (gray region).

### Biophysical groups do not generally map to sequence similarity

Biophysical groups are generally not concordant with sequence-similarity based groups used to infer shared protein function using traditional bioinformatics approaches (Figs. 3c, 3g, S7 and S8). Exceptions include three small PBD families: 14-3-3 (Fig. 3f), WH1, and SWIB (Fig. S7). For the 14-3-3 family, this finding is consistent with a previous experimental study that demonstrated that affinity differences between the seven human isoforms strongly correlate with their sequence divergences. ^18^ Nevertheless, their affinity differences cannot be traced back to amino acid changes at specific positions within their highly conserved all-helical structure, including within their phosphopeptide binding pocket, which is in fact identical across isoforms. ^18^ This suggests that QSM models how combinations of amino acid differences spanning the entire 14-3-3 fold impact binding to yield biophysical group assignments that capture experimentally determined affinity differences. Indeed, the two biophysical groups distinguish the *ω* and *ε* isoforms (Group P, pink in Fig. 3g) from the other four (Group B, blue) to recapitulate published groupings based on (i) the number of shared protein interaction partners detected in two cell lines with affinity-purification mass-spectrometry (AP-MS) and (ii) average binding strength differences for peptides derived from E6 oncoproteins of tumorigenic human papillomaviruses measured using *in vitro* assays. ^18^

### Biophysical groups vary in a!nity and selectivity

Biophysical groups capture differences in domain binding strength and selectivity in addition to sequence-specific binding preferences for particular peptidic sites in the human proteome. Indeed, biophysical group membership is apparent when domains are projected onto a lower dimensional space constructed using principal component analysis to preserve most of the variance in their affinities for human peptidic sites (Fig. 3d and S9). We find that much of the variance in their affinities reflects differences in binding strength, which we quantify by each domain’s affinity for its most favored 2% of peptidic sites to guard against incorrect binder classifications made at a 1% false discovery rate (FDR) (as indicated on SHANK3’s affinity distribution in Fig. 3a): The first principal component largely aligns with changes in binding strength observed within most PBD families (Fig. 3e and S9). At the strong end of the binding strength spectrum are domains from SWIB, FERM, Ankyrin, ARM, WDR, SH3, and WW families, which have predicted nanomolar binding strengths; at the weak end are domains from 14-3-3 and PTB families, which have predicted upper micromolar to low millimolar strengths (we note that very weak binding affinities may indicate false positive predictions). Nevertheless, most domains have modest, micromolar binding strengths (Fig. 3e, f and S9).

We find that a smaller proportion of the variance generally reflects differences in selectivity—how much stronger each domain binds its most favored sites relative to the pool of available binding sites in the proteome—which we quantify by each domain’s median affinity relative to its binding strength (as indicated on DLG2-2’s affinity distribution in Fig. 3a): The second principal component largely aligns with changes in selectivity observed within many PBD families (Fig. 3e and S9). Domains from the SH3, WW, Chromo, Bromo, PTB, TK, and PTP families, many of which recognize PTMs, exhibit the highest selectivities, in some instances favoring specific peptidic sites 40-60 fold more than those recognized by other members of the same family (Fig. 3e, f and S9). The majority of domains, though, have modest selectivity, showing no more than a 10 fold preference for specific sites. PDZ is one example of such promiscuous PBD families, which divide into relatively few biophysical groups compared to the total number of domains in the family (Fig. 3b, g). By contrast, selective families, such as SH3, split into many biophysical groups with distinctive binding preferences (Fig. 3g).

The observation that binding strength and selectivity each generally align with different, orthogonal principal components suggest that these two biophysical properties are largely decoupled. Indeed, pairs of domains from the same PBD family may have similar selectivities but order-of-magnitude different binding strengths, such as seen when comparing PDZs belonging to Group B (blue) to those in Group Y (yellow) in Fig. 3e. Yet, in other cases, both selectivity and strength may increase for one domain relative to another, such as seen when comparing SH3s belonging to Group B to those in Group G (green) in Fig. 3e. Overall, this suggests that binding strength and selectivity can be independently tuned through subtle changes in PBD sequence to sample the landscape of possible binding profiles.

### Experimental validation of PDZ assignments to biophysical groups

To assess the quality of our classification system, we experimentally compared the binding affinities of PDZ domains across biophysical equivalence groups. Specifically, we assayed two PDZ domains from Group B, the second PDZ domains of disk large homologue 1 and 2 (DLG1-2 and DLG2-2), and one PDZ domain from Group G, that of SHANK3, against a panel of 12 peptides derived from the human proteome whose affinities to all three PDZs were predicted with high confidence (Fig. 3h). Splitting the panel between peptides with predicted 10-fold stronger affinities for SHANK3 over either DLG1/2-2 and those predicted to show the inverse relationship allowed us to test if QSM correctly quantifies both differences and similarities of domains’ binding profiles. Additionally, this panel included thee peptides with published *K*_D_’s included in QSM’s training data so that we could quantify experimental error due to variation in assay conditions and methods used by different labs. None of the other peptides were included in the training set. Using a fluorescence polarization (FP) assay, we determine *K*_D_’s that differ from those published by a factor of 3 on average and by roughly an order of magnitude for 2 out of 5 interactions (Fig. 3h, orange points, peptides 1 and 9), setting a standard for determining if QSM predictions match experimental accuracy.

QSM predictions generally agree with our experimentally determined *K*_D_’s for these interactions within a factor of 2-5, with 83% agreeing within an order of magnitude, further demonstrating that QSM quantifies affinities with comparable accuracy to experimental methods (Fig. 3h and Table S2). While *K*_D_’s could not be reliably quantified from our assays for the weakest interactions, we were able to quantify affinities for each peptide to at least one PDZ domain, and both DLG1-2 and DLG2-2 if one was reliably determined. These measurements confirm that DGL1-2 and DLG2-2 have highly similar binding profiles, with their affinities for the same peptide differing by only 0.29 kcal/mol on average (nearly matching a predicted average difference of 0.23 kcal/mol), supporting our classification of them into the same biophysical equivalence group. These measurements also confirm that SHANK3’s binding preferences are distinct from DLG1/2-2, with QSM correctly identifying which domain binds 11 of the 12 peptides more strongly. Consistent with the prediction that SHANK3 is a more promiscuous binder than DLG2-2 (Fig. 3a, affinity distributions), we were able to quantify the affinities of most of the peptide panel (75%) for SHANK3 but fewer (58%) for DLG1/2-2. Overall, these experiments support our classification of these domains into their respective biophysical groups.

Moreover, these results demonstrate that QSM captures differences in selectivity even for domains with similar consensus binding motifs. Both DLG1/2-2 and SHANK3 are classified in the literature as class I PDZ domains that recognize c-terminal motifs of the type X[T/S]XΦ, where X is any amino acid and Φ is hydrophobic. ^54^ Although all peptides in our panel display this motif except peptide 4 and only peptide 5 displays the XSXΦ variant, they bind members of the two biophysical groups with quantitatively distinct affinities (Fig. 3h). Thus, established consensus motifs constructed from high-throughput qualitative readouts or small scale mutational studies of an individual motif derived from a single protein interactor insufficiently characterize PBD selectivity. Our data also indicates that residues outside the core motif must be considered to explain affinity differences, further mirroring findings in ASHI ^9^ and recent deep mutational scans of six PBD–peptide interactions. ^56^ Biophysical equivalence groups constructed from proteome-wide QSM predictions provide a modern classification system without such limitations.

### QSM generalizes to multivalent protein interactions

Because many human proteins contain multiple PBDs and/or peptidic sites, their observed binding preferences reflect the combined preferences of their constituent binding modules, ^57^ which can combine in cooperative and competitive ways. To model such multivalent protein–protein interactions, we use the QSM free energy function to parameterize statistical mechanical models of modular proteins that form both intra- and inter-molecular PBD–peptide interactions: We compute the energies of all possible bound and unbound conformations to estimate PPI affinities from ratios of partition functions (Fig. 4a). We refer to these models as QSM for proteins (QSM/P). QSM/P explicitly models multivalent binding and local steric constraints imposed on adjacent peptidic sites and intramolecular interactions between PBDs and peptidic sites that overlap in sequence. Thus, QSM/P accounts for increases in PPI affinity due to avidity and for biases in proteins’ conformational landscapes due to inter-PBD/peptide competition.

**Fig. 4.**
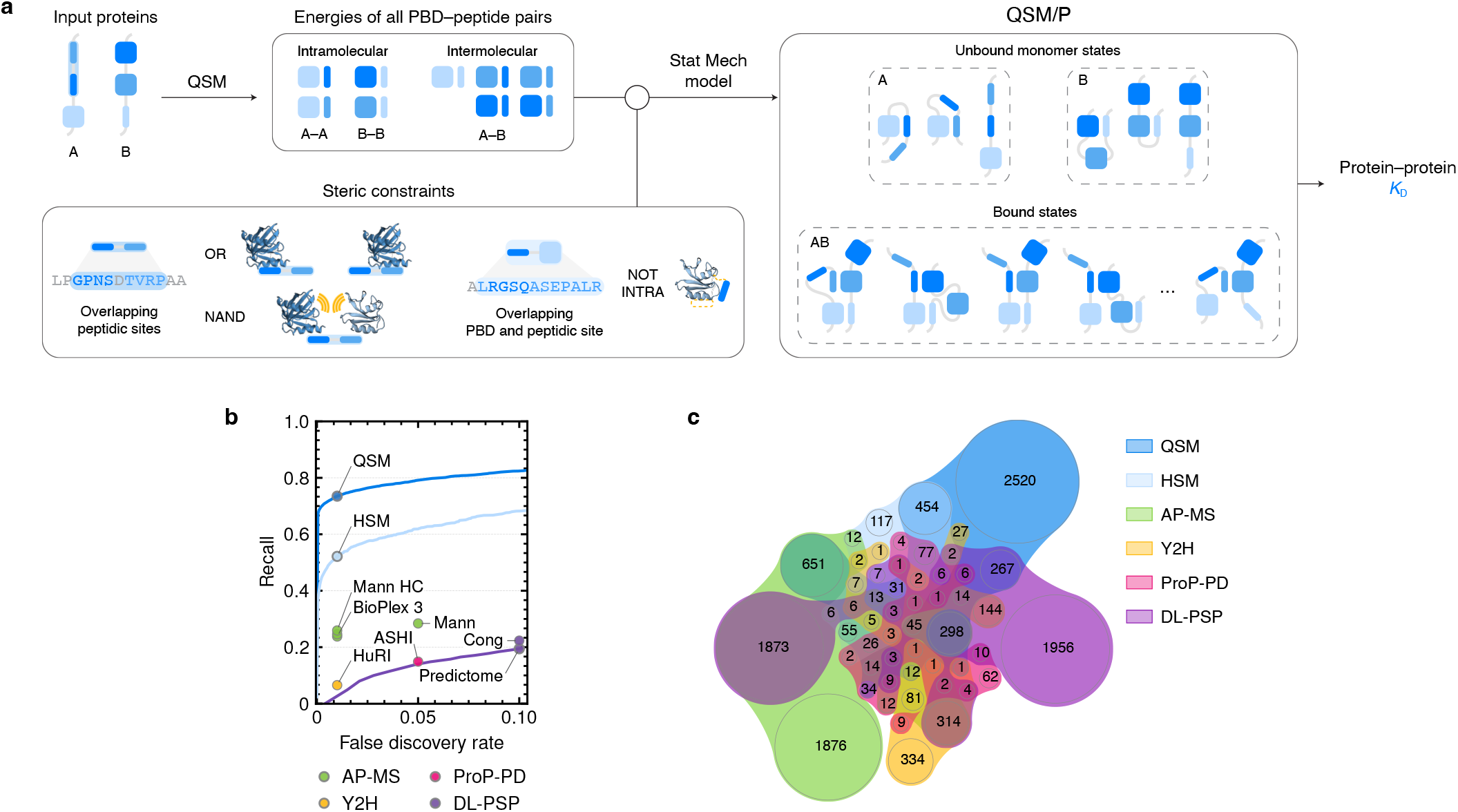
QSM/P models multivalent protein–protein interaction mechanisms. **a.** Interactions between modular proteins are modeled with QSM/P by first decomposing proteins into all possible domain–peptide interactions and estimating their interaction energies with QSM, and then computing the *K*_D_ of the PPI from the ratio of unbound to bound state partition functions. Energies of each unbound monomer and bound dimer conformation are computed as the sum of bound domain–peptide interactions, and only conformations that respect steric constraints imposed on peptidic sites and/or PBDs overlapping in protein sequence contribute to the partition function. **b.** Recall versus FDR of PPIs confirmed by multiple low-throughput sources in HINT ^58^for QSM, previous state-of-the-art model HSM ^29^, a!nity-purification mass-spectrometry (AP-MS) datasets (BioPlex 3 for HEK293T and HCT116 cell lines, ^2^ Mann HC, and Mann ^59^), yeast-two hybrid (Y2H) dataset (HuRI ^1^), proteomic peptide phage display (ProP-PD) dataset (ASHI ^9^), and deep learning protein structure prediction (DL-PSP) datasets (Cong ^4^ and Predictome ^60^). FDRs taken from associated publications. Curves plotted when recall could be evaluated at multiple FDRs from a single dataset. **c.** Generalized quasi-proportional Venn diagram ^61^ of PPIs recalled by each method at FDR indicated in **b**.

To evaluate the accuracy of QSM/P, we assessed its recall of protein interactions confirmed by multiple independent sources in HINT ^58^using low-throughput assays. We compared it to (i) high-throughput experimental methods developed to broadly detect PPIs (AP-MS ^2,59^and yeast two-hybrid (Y2H) ^1^) and to specifically detect PBD/motif-mediated interactions (proteomic peptide phage display (ProP-PD) ^9^) and (ii) computational screening approaches developed to map interactomes using deep learning protein structure prediction (DL-PSP) methods, namely AlphaFold and RF2-PPI. ^4,60^To fairly compare methods, recall was computed on the subset of PPIs detectable by each (*e.g.*, only PPIs involving the baits used in AP-MS or ProP-PD). At equivalent FDRs, QSM/P recalls roughly three times more PPIs than the best high-throughput method—experimental or computational—and outperforms our previous model of PBD-mediated PPIs, HSM/P, ^29^which is limited to fewer PBD families and binding chemistries and does not quantify affinities (Fig 4b).

Each method recalls a distinct but overlapping set of PPIs that reflect its biases towards given types of interactions (Fig 4c). AP-MS, Y2H, and DL-PSP methods generally detect stronger, globular, and well-structured PPIs, whereas these types of interactions remain blind spots for ProP-PD and QSM/P, which specifically target weak, PBD-mediated interactions. Of the HINT PPIs that can be modeled with QSM/P, 91% of those experimentally detected with ProP-PD in ASHI ^9^at a 5% FDR are also recalled by QSM/P at a 1% FDR. This high concordance demonstrates that QSM/P achieves experimental accuracy on PBD-mediated PPIs and suggests that QSM/P can be integrated into experimental screening pipelines to yield larger, higher-confidence sets of PPIs. QSM/P is, thus, ideal for mapping the PBD-mediated interactome and discovering new PPIs that challenge existing methods.

### QSM infers the physical organization of signaling networks

Applied proteome-wide, QSM/P predicts roughly two million PPIs with submillimolar affinities at a 1% FDR. While QSM/P predicts physically plausible interactions, additional cellular mechanisms determine which protein interactions occur in a particular cellular context. Notably, proteins must be colocalized within cellular subcompartments to physically interact. Since this is not explicitly modeled by QSM/P, we further filter our predictions to protein pairs with experimental evidence of overlapping subcellular localization distributions from the Cell Atlas, part of the Human Protein Atlas (HPA). ^62^We refer to the resulting network as QSM/N (Fig. 5a). Subcellular distributions have been determined for 85% of human proteins, and this imposes a limit on the maximum number of proteins included in QSM/N. QSM/N includes 455,384 interactions between 11,298 proteins (55% of human proteins), covering a significant fraction of the potential interactome, which is estimated to contain over 1 million PPIs. ^1^The scale of QSM/N exceeds that of current experimental maps, which cover roughly 50,000 to 120,000 PPIs in HuRI ^1^and BioPlex, ^2^respectively, and expands the number of PBD-mediated PPIs uncovered in ASHI ^9^by over 25-fold.

**Fig. 5.**
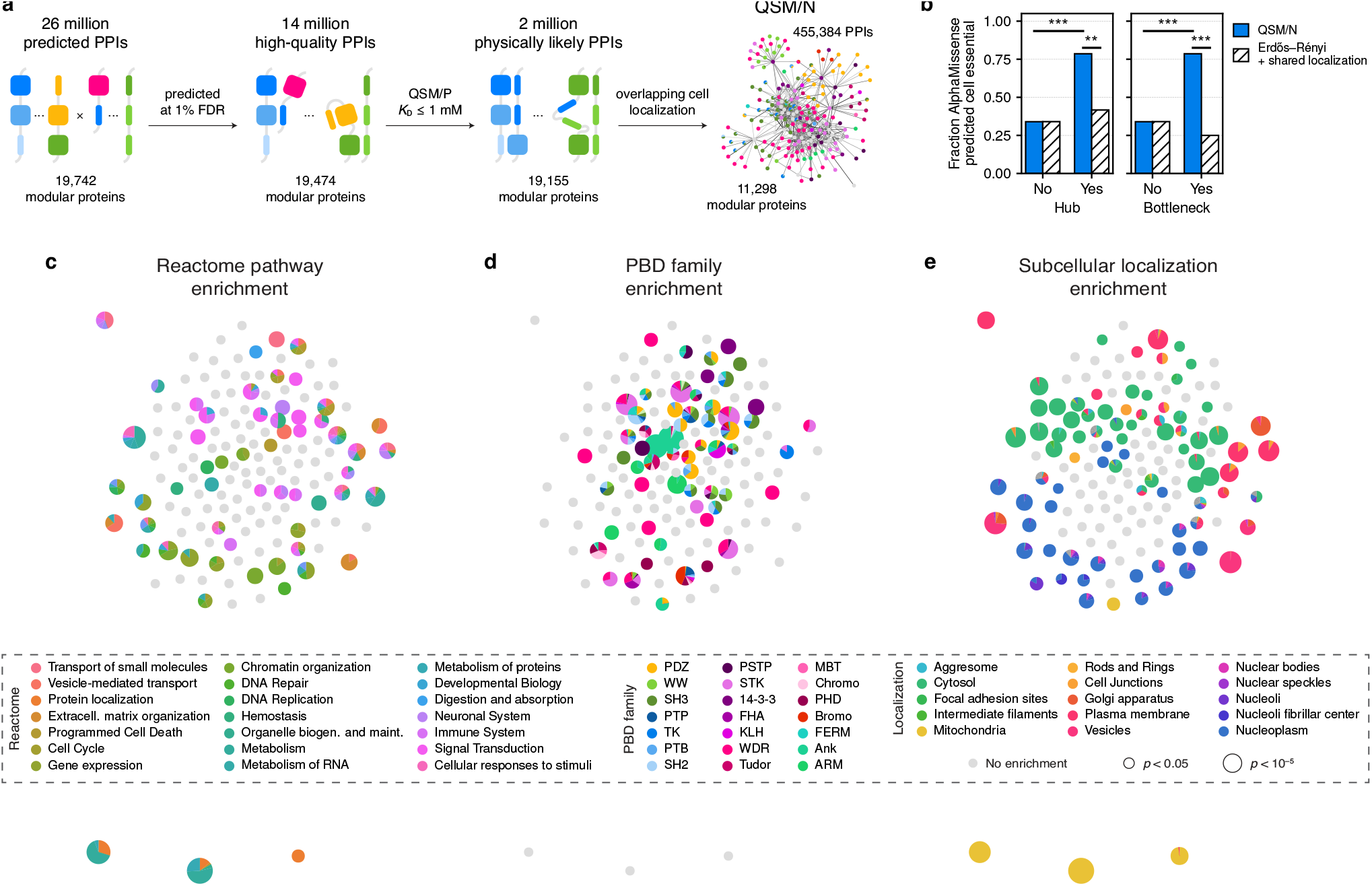
Functional modules emerge in predicted PBD-mediated protein interaction network. **a.** Pipeline to construct QSM/N. QSM/N subnetwork for mitogen-activated protein kinase (MAPK) signaling with nodes corresponding to proteins (colored by PBD family composition) and edges to QSM/P predicted *K*_D_’s (stronger interactions are darker) laid out using force-directed layout. ^63,64^ **b.** Protein hubs, distinguished by largest closeness centrality, and bottlenecks, distinguished by largest betweenness centrality, in QSM/N are significantly enriched in AlphaMissense predicted cell essential proteins over non-hub and non-bottleneck proteins and proteins with the highest centralities in a null, Erd?os-Rényi network model that satisfies the constraint of shared subcellular localization used to construct QSM/N (*** *p<* 0.001, ** *p<* 0.01, binomial test). **c-e.** Coarsened QSM/N with nodes corresponding to protein modules and laid out using a force-directed layout such that modules with similar connectivity patterns cluster together illustrates large-scale network structure. Edges between modules not shown. Each node is represented by a pie chart that denotes the enrichment score (log *p*, corrected for multiple hypothesis testing with the Benjamini-Hochberg method) of (**c**) Reactome ^65^biological processes, (**d**) subcellular localization, and (**e**) PBD family composition of proteins assigned to each module. Modules with fewer than four proteins or without any statistically significant enrichment (*p* ≥ 0.05) colored gray. Nodes sized according to the significance of the most enriched category. Categories with dominant enrichment values for at least one module listed.

Given that QSM/N comprehensively and unbiasedly maps the PBD-mediated interactome—without systematic methodological biases towards strong, evolutionary conserved, or well-structured interactions or highly expressed proteins that shape interactomes mapped with other methods ^1,2,4,60^—we hypothesized that it captures molecular mechanisms underlying cell function. To test this hypothesis, we examined if biology that has been previously uncovered in experimentally mapped protein interaction networks can be rediscovered in QSM/N.

The topology of experimental protein interaction networks can inform on protein function. ^66^Proteins that act as hubs— highly connected proteins with high closeness centrality—and bottlenecks—critical points of information flow corresponding to proteins with high betweenness centrality—tend to be essential or disease-associated. ^67–69^Consistent with this observation, we find that hubs and bottlenecks in QSM/N are significantly enriched for proteins that are predicted to be essential by AlphaMissense ^70^(*p <* 0.001, binomial test) (Fig. 5b). Additionally, randomly rewiring QSM/N under that constraint that new edges only link proteins with evidence of overlapping subcellular localization distributions—the only explicit experimental information used to construct QSM/N—significantly abolishes this enrichment (*p <* 0.01 for hubs and *p <* 0.001 for bottlenecks, binomial test), demonstrating that QSM/P predictions are necessary to construct a network that reflects protein function.

Interaction networks, furthermore, contain information about how collections of proteins coordinate more complex cellular functions, such as metabolism, growth, and differentiation. Proteins that share many interaction partners in the network likely play similar functional roles within a cellular pathway, and communities of interacting proteins can correspond to assemblies that perform specific, higher-order functions. ^1,2,71,72^This motivated us to next ask if QSM/N organizes into modules with known cellular functions. We first partition QSM/N into modules of proteins with similar binding strengths to all others by fitting a nested stochastic block model (SBM), ^73,74^specifically the degree corrected, weighted variant, ^75^to QSM/N (Fig. S10). As nonparametric generative models for uncovering network structure via statistical inference, nested SBMs make minimal assumptions about the topological features that delineate modules, resolve small modules even in large networks, and avoid identifying spurious modules resulting from random noise. ^76^ By taking this unbiased approach to community detection, we find both assortative and disassortative modules of proteins (Fig. S11) among the 149 identified in QSM/N. We then examined each of the 104 modules with at least four proteins (Fig. S12) for enrichment in cellular pathways cataloged in the Reactome database. ^65^ The majority of modules (57%) map to known biological processes (5% Benjamini-Hochberg corrected FDR) spanning nearly all core pathways: metabolism, gene regulation and expression, protein localization and molecular trafficking, cellular responses to stimuli and signal transduction, and cell cycle and apoptosis are some of the most significantly enriched (Fig. 5c). Thus, QSM/N enables the (re)discovery of complex biological processes mediated by protein interactions.

Remarkably, these cellular-scale pathways emerge from molecular-scale protein interactions predicted from sequence information alone. We can leverage this aspect of QSM/N to gain fundamental insights into the physical mechanisms underlying cellular function—insights that remain elusive when enrichment analysis is similarly applied to experimental transcriptomic or proteomic data to identify genes that collectively regulate biological pathways. ^76^We can examine physical organization at two levels within QSM/N: at the level of cellular compartments, which captures micron-scale structure, and at the level of molecular interactions, which reflects nanometer-scale structure. QSM/N modules enriched for the same cellular processes cluster together as a result of mechanisms operational at both of these physical scales (Fig. 5c-d). Most modules with significant enrichment for a known biological process are also enriched for particular subcellular localizations (86%) (Fig. 5c, e, S13, and S14), typically the organelles and cellular structures whose textbook function matches the module’s inferred function. Many QSM/N modules, including those enriched for distinct functions, map to the same subcellular compartments; proteins in these modules are distinguished not by different subcellular localizations but by different molecular recognition mechanisms and binding profiles. The majority of functional modules are also enriched for specific PBD families (58%) (Fig. 5c, d, and S15) and peptidic chemistries (63%) (Fig. S16), and eight of the nine functional modules lacking a clear subcellular localization are instead enriched for specific PBD families. Thus, proteins’ molecular binding preferences and subcellular localizations, combined, physically organize the human PBD-mediated interactome into functional modules.

## Discussion

QSM quantitatively predicts PBD–peptide interaction affinities with experimental-level accuracies at an unprecedented scale, realizing the potential of deep learning to provide a link from protein sequence to binding affinity. QSM achieves this by transforming a problem previously considered unsolvable due to data limitations into one learnable through data harmonization as a direct consequence of statistical mechanics. Structuring learning within a Single Objective, Multiple Data framework provides the flexibility to train models on larger datasets synthesized from both low-throughput affinities and high-throughput binary measurements and can foreseeably advance quantitative modeling of biomolecular interactions more generally. Combined with a biophysically informed architecture that facilitates knowledge transfer across biochemically and structurally heterogeneous PBD families with varying degrees of experimental characterization (Fig. S4), QSM triples the number of PBD families modeled with high fidelity to-date, predicts the effects of human genetic variation on SLiM-mediated interactions, and shows promising ability to generalize to chaperones, RNA-binding domains, and enzymes whose peptide recognition capabilities have been detected experimentally only very recently. ^9^

A key feature of QSM is the prediction of absolute thermodynamic constants with estimates of their own prediction error. Quantifying PBD–peptide dissociation constants allows us to universally compare their binding preferences and enables us to build models of increasing complexity and scale—spanning modular proteins to proteome-scale interaction networks—from these predictions. Leveraging QSM’s self-estimates of prediction accuracy in both these endeavors provides confidence in the biophysical insights and novel interactions that we uncover from applying QSM proteome-wide. Quantifying the binding preferences of thousands of human PBDs reveals recurring biophysical archetypes with characteristic binding strengths and selectivities that are largely decoupled within a PBD family—an observation that suggests these two properties can be programmed independently through sequence engineering to create PBD-based biosensors ^77,78^ and protein therapies ^79,80^ with target affinity-specificity profiles. We distill these findings into a classification system of biophysical equivalence groups, which comprise sets of binding modules with similar binding preferences that are reused and combined within different proteins to create more complex recognition mechanisms. This repertoire of molecular recognition mechanisms, combined with differences in protein subcellular localization, gives rise to a hierarchical network structure that organizes proteins into functional pathways and emerges from applying QSM at proteome scale.

The approach taken by QSM marks a significant advance in our ability to model and probe PBD-mediated signaling and regulatory networks: quantitative models of protein interaction circuits provide a means to predict and explain the impact of disease mutations and altered protein expression levels on cellular behavior; ^81^ proteome-scale models offer opportunities to uncover points of pathway crosstalk ^7^ and reveal convergent mechanisms of cellular information processing. ^82^ Expanding QSM models to additional PBD families can be done efficiently through active learning, ^83,84^ leveraging QSM’s uncertainty estimates to guide experimental data acquisition. Combining QSM with protein structure predictors ^3,25,43^and coarse-grained models of intrinsically disordered protein regions ^85,86^ can structurally resolve multivalent PPI ensembles to better account for physical mechanisms underlying avidity ^87^and cooperativity. ^88^ Incorporating complementary sources of experimental data in addition to subcellular localization measurements, for example, from time-resolved phosphoproteomics ^89^ or spatial transcriptomics ^90^ would further enable modeling of signaling dynamics and cell-cell communication. Our work thus establishes an extensible computational framework to construct mechanistic models of cell signaling and a conceptional foundation upon which to build a comprehensive understanding of biological computation.

## Code and data availability

Release of all code and datasets is in preparation.

## Acknowledgments

J.R.R. acknowledges a Burroughs Wellcome Fund Career Award at the Scientific Interface (Award No. 1361252) and Fellowship of The Jane Coffin Childs Memorial Fund for Medical Research. This work was supported by National Science Foundation grant 2441001 to N.H.S. L.P.C. was supported by a National Science Foundation Graduate Research Fellowship.

## Notes

### Competing Interest Statement

M.A. is a member of the scientific advisory boards of Cyrus Biotechnology, Deep Forest Sciences, Nabla Bio, Oracle Therapeutics, and Achira.

