## Supplementary Information for "Quantitative machine learning of protein interactions reveals the multiscale organization and molecular syntax of signaling networks"

### Supplementary Tables

**Table S1 Ablations demonstrate QSM learns effectively from heterogenous data using multiple mechanisms**. Intra-family performance of QSM, QSM/D, QSM/ID, and ablated QSM/ID models evaluated in 5-fold cross-validation on quantifying affinities ($\log K_\mathrm{D}$) and classifying binders.

| Model | Median intra-family | | |
|---|---|---|---|
| | MAE | Pearson $R$ | AUROC |
| QSM | 0.53 | 0.67 | 0.96 |
| QSM/D | 0.65 | 0.63 | 0.94 |
| QSM/ID | 0.56 | 0.58 | 0.91 |
| no pLM | 0.65 | 0.49 | 0.86 |
| $K_\mathrm{D}$ as binary data | 1.38 | 0.06 | 0.52 |
| $K_\mathrm{D}$ data | 0.67 | 0.44 | 0.55 |
| binary data | 1.04 | 0.36 | 0.94 |

**Table S2 Predicted and experimentally measured affinities of PDZ domains.**

| Peptide | DLG1-2 $K_D$ (micromolar) | | | DLG2-2 $K_D$ (micromolar) | | | SHANK3 $K_D$ (micromolar) | | |
|---|---|---|---|---|---|---|---|---|---|
| | QSM | FP | Literature [51] | QSM | FP | Literature [51] | QSM | FP | Literature [51] |
| AHFSSLESEV | $3.48 \pm 1.02$ | $7.70 \pm 1.04$ | — | $4.48 \pm 1.43$ | $4.05 \pm 0.74$ | — | $63.83 \pm 28.06$ | $72.51 \pm 10.01$ | — |
| PQAQAENTAF | $147.09 \pm 50.93$ | N.D.[a] | — | $234.90 \pm 84.68$ | $969.80 \pm 31.60^b$ | — | $8.20 \pm 3.78$ | $6.50 \pm 0.67$ | — |
| QIAGTKSTTV | $7.00 \pm 3.03$ | $15.21 \pm 7.37$ | — | $12.18 \pm 4.92$ | $5.95 \pm 0.52$ | — | $278.06 \pm 131.66$ | $120.70 \pm 29.17$ | — |
| TTVRCLEAEV | $7.27 \pm 2.79$ | $438.37 \pm 392.90$ | — | $11.66 \pm 4.81$ | $477.20 \pm 305.60$ | — | $223.73 \pm 108.52$ | N.D.[a] | — |
| STLTIFETAL | $61.68 \pm 18.86$ | $330.30 \pm 203.50$ | 27.10 | $86.34 \pm 25.11$ | $65.73 \pm 4.38$ | 71.05 | $9.24 \pm 3.94$ | $0.33 \pm 0.15$ | 2.86 |
| SHSSKGETAV | $16.52 \pm 4.63$ | $16.59 \pm 4.13$ | 16.41 | $28.25 \pm 7.64$ | $17.99 \pm 6.22$ | 28.07 | $328.12 \pm 128.53$ | $431.20 \pm 56.92^b$ | — |
| EPPEEFDTAL | $88.35 \pm 31.66$ | $518.60 \pm 238.60^b$ | — | $148.25 \pm 49.91$ | $791.70 \pm 42.67^b$ | — | $3.06 \pm 1.57$ | $11.53 \pm 3.17$ | — |
| NQLAWFDTDL | $217.45 \pm 74.55$ | $771.70 \pm 317.50^b$ | — | $242.88 \pm 80.59$ | $835.60 \pm 288.80^b$ | — | $17.95 \pm 9.74$ | $83.31 \pm 35.63$ | — |
| DFREDDDTAL | $335.33 \pm 100.18$ | $564.90 \pm 137.30^b$ | — | $395.46 \pm 109.81$ | $342.07 \pm 62.35^b$ | — | $18.28 \pm 9.44$ | $18.00 \pm 6.51$ | — |
| SGSLKVMTTV | $3.51 \pm 1.32$ | $14.78 \pm 8.89$ | — | $4.55 \pm 1.67$ | $22.97 \pm 21.06$ | — | $256.19 \pm 107.91$ | $922.80 \pm 959.10^b$ | — |
| TRVVDQITTV | $9.53 \pm 3.63$ | $157.00 \pm 59.20$ | — | $20.46 \pm 7.02$ | $152.90 \pm 47.28$ | — | $188.66 \pm 90.84$ | $40.23 \pm 5.97$ | — |
| LRKPEADTAL | $123.85 \pm 42.94$ | N.D.[a] | — | $217.09 \pm 65.29$ | $632.60 \pm 467.20^b$ | — | $8.08 \pm 4.55$ | $34.02 \pm 16.18$ | — |

[a] Not determined.

[b] Estimated $K_D$. Too weak to reliably quantify.

### Supplementary Figures

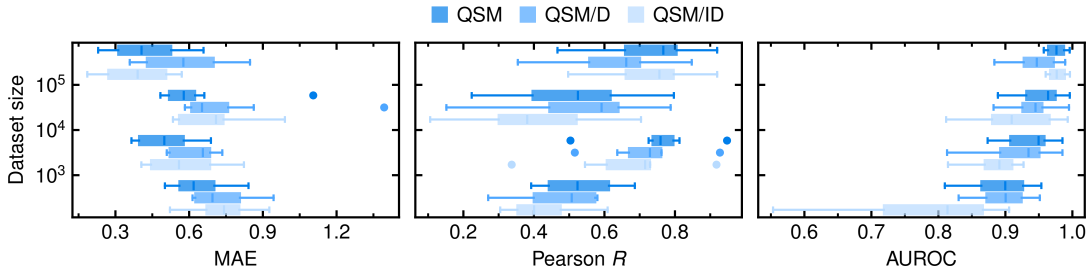


**Fig. S1 Model ensembling mitigates PBD family dataset size disparities.** Performance of QSM/ID, QSM/D, and QSM models for each PBD family stratified by training dataset size.

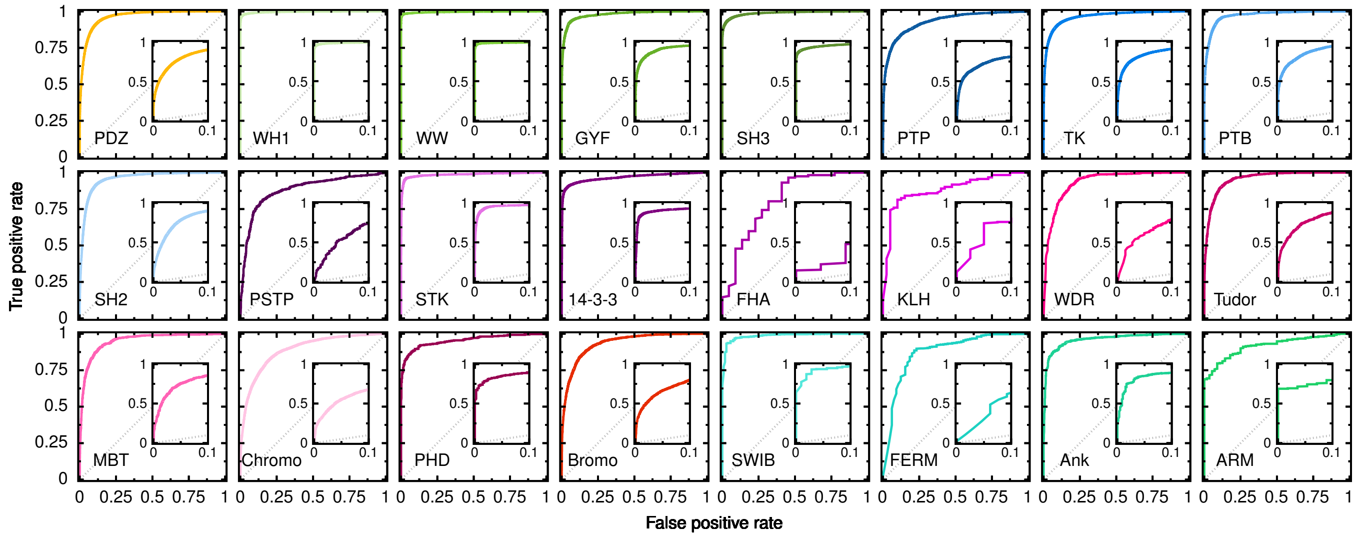


**Fig. S2 QSM accurately classifies PBD–peptide interactions.** Receiver operating characteristic curves for each in-distribution PBD family. Evaluations performed in 5-fold cross-validation, in which unique PBD–peptide pairs were randomly assigned to each fold.

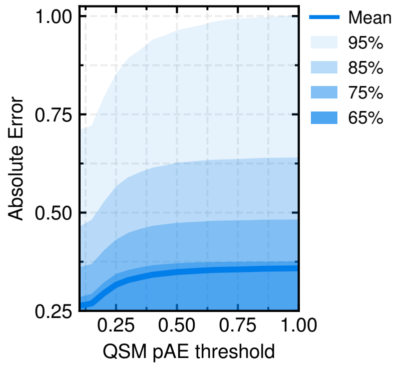


**Fig. S3 High confidence predictions correspond to QSM pAE of 0.25.** True absolute error as a function of QSM predicted absolute error (pAE) threshold. Mean and percentile true absolute errors from 5-fold cross-validation on randomly split PBD–peptide pairs plotted.

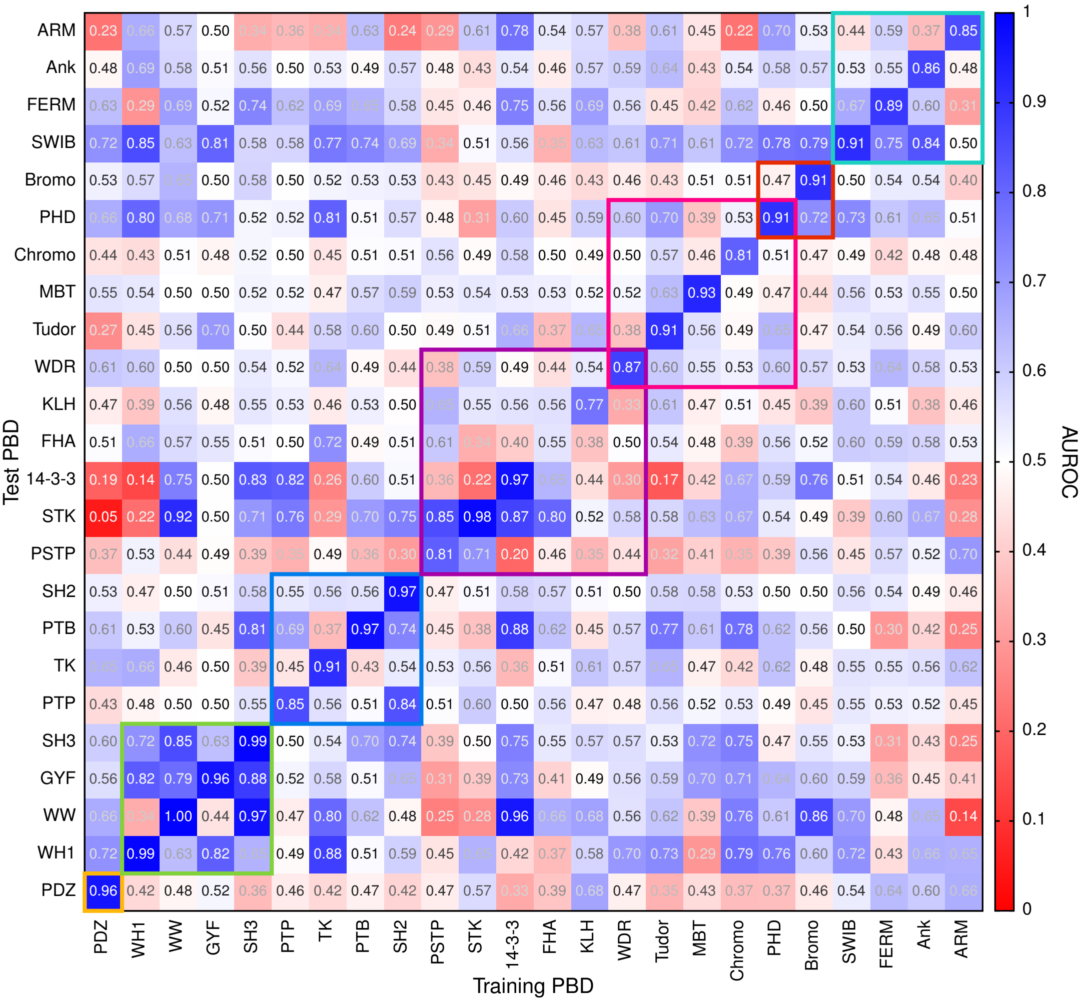


**Fig. S4 QSM can classify interactions mediated by out-of-distribution PBD families.** Classification performance of QSM/ID models trained using 5-fold cross-validation on data from one PBD family and tested on all other families. Diagonal entries report intra-family AUROC aggregated over all folds. Off-diagonal entries report family-transfer AUROC averaged over all folds. Colored squares outline PBD families with shared binding chemistries (orange for c-terminal, green for polyproline, blue for phosphotyrosine, purple for phosphoserine/threonine, pink for methyllysine/arginine, red for acetyllysine, and teal for other, mostly hydrophobic peptidic motifs).

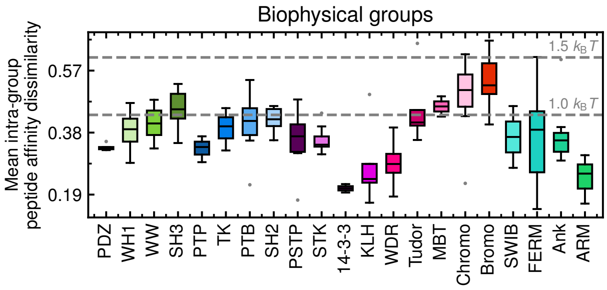


**Fig. S5 Biophysical groups cluster PBDs together if their predicted binding affinities for peptidic sites across the proteome are roughly indistinguishable from thermal energy fluctuations.** Mean intra-group similarity of peptide affinities. For domains $A$ and $B$, the similarity of their affinities for the $N$ peptidic sites in the human proteome is calculated as $\left(N \sum_i^N \log\left(K_{\mathrm{D}_i}^{(A)}/K_{\mathrm{D}_i}^{(B)}\right)^2\right)^{1/2}$.

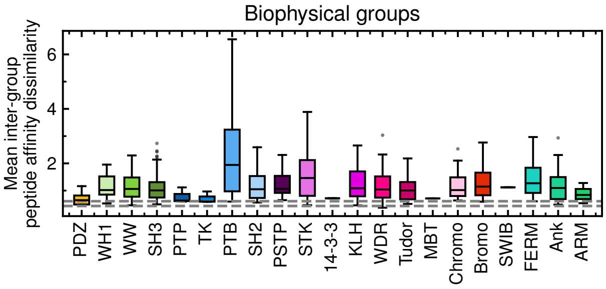


**Fig. S6 Biophysical groups separate PBDs if their predicted binding affinities for peptidic sites across the proteome differ by an order-of-magnitude.** Mean inter-group similarity of peptide affinities. For domains $A$ and $B$, the similarity of their affinities for the $N$ peptidic sites in the human proteome is calculated as $\left(N \sum_i^N \log\left(K_{\mathrm{D}_i}^{(A)}/K_{\mathrm{D}_i}^{(B)}\right)^2\right)^{1/2}$. Dashed lines indicate 1 and $1.5k_{\mathrm{B}}T$.

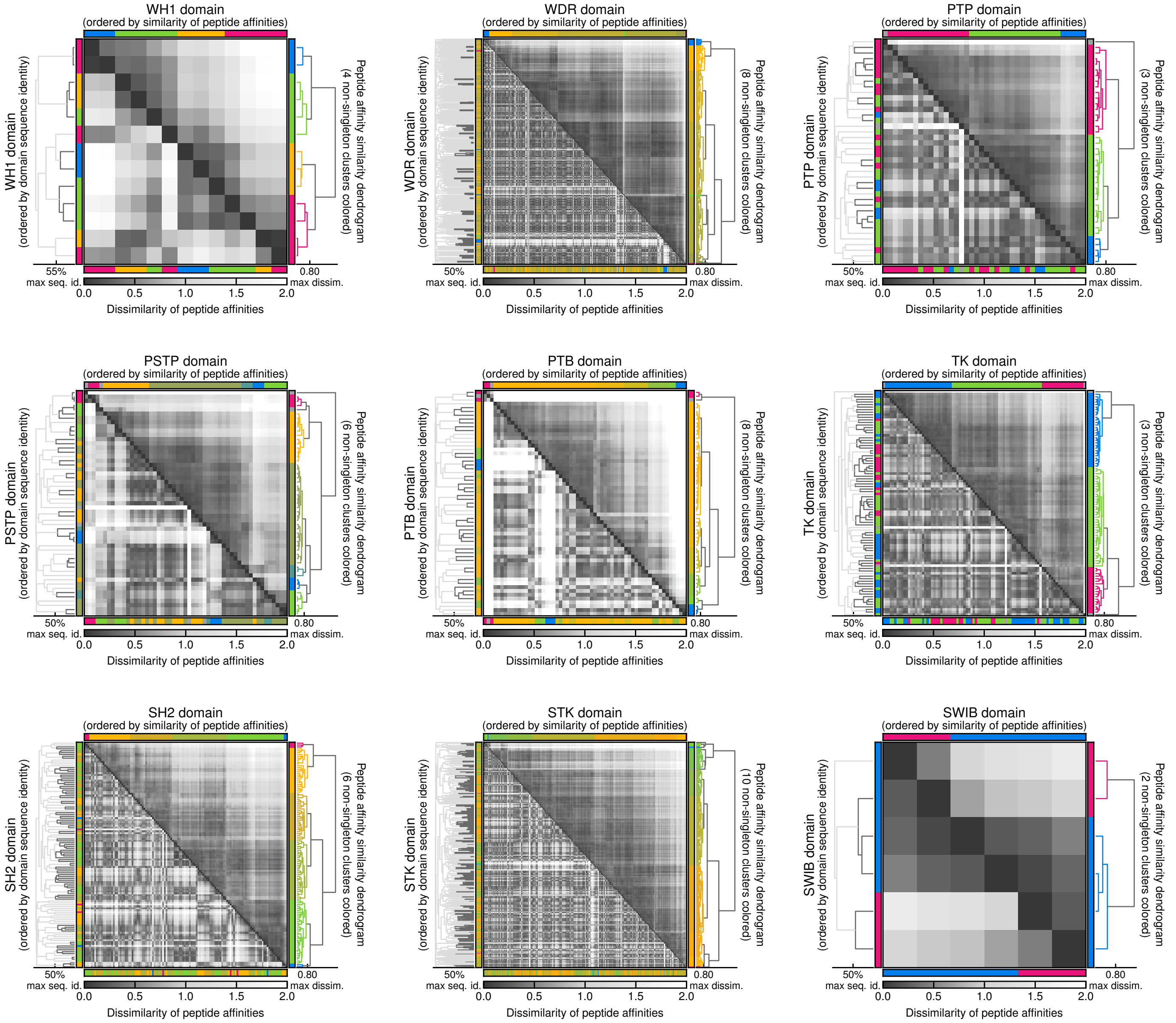


**Fig. S7 Proteome-wide QSM affinity predictions enable classification of PBDs into biophysical equivalence groups.** Similarity matrices of peptide affinities. Upper half is ordered by domains' similarity of peptide affinities, and lower half is ordered by their sequence similarities. Colors indicate biophysical group assignments and are assigned based on cophenetic distance. Clustering thresholds indicated on each dendrogram. Domains that cluster together based on sequence identity colored dark gray on dendrogram.

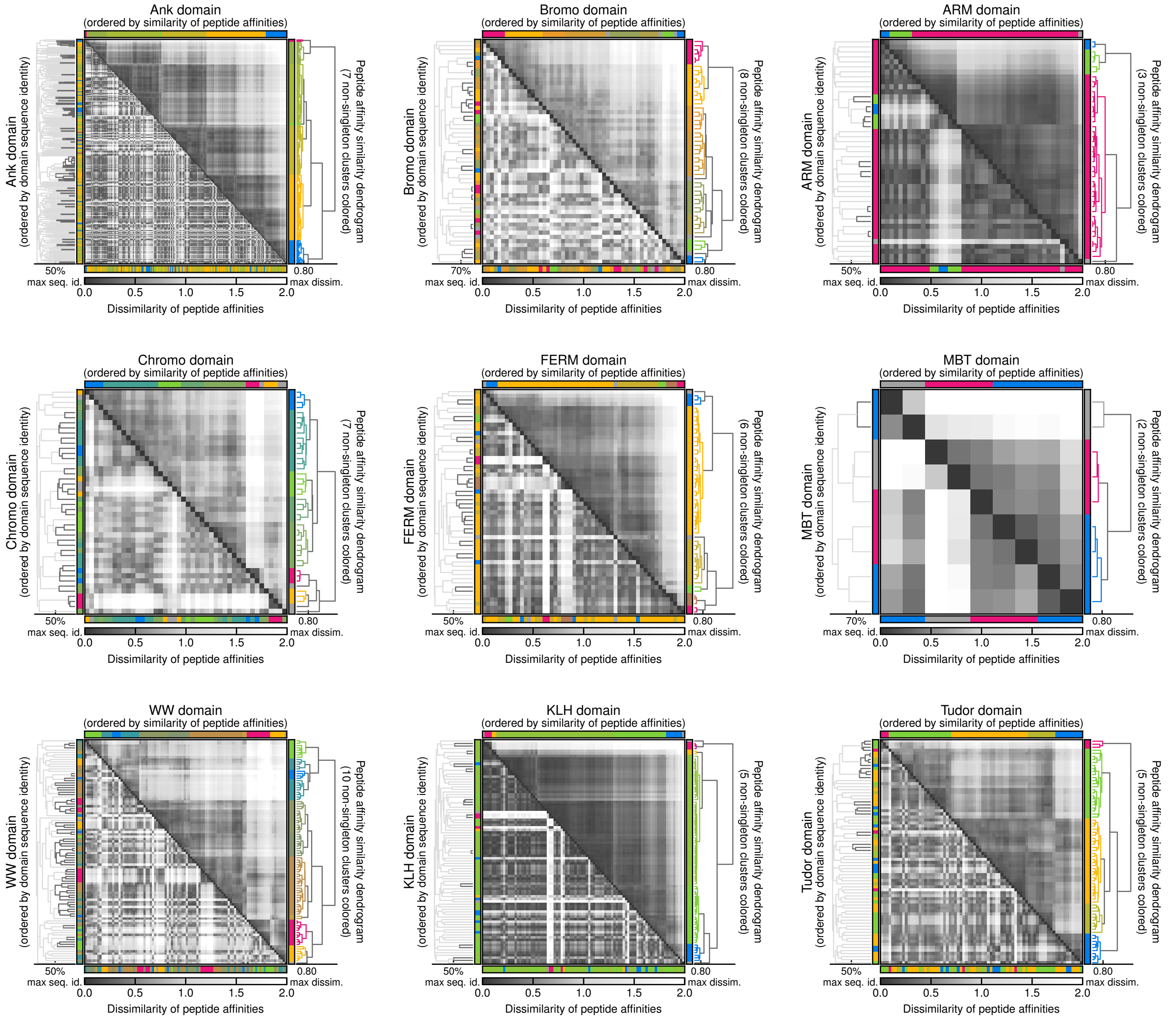


**Fig. S8 Proteome-wide QSM affinity predictions enable classification of PBDs into biophysical equivalence groups (continued).** Similarity matrices of peptide affinities. Upper half is ordered by domains' similarity of peptide affinities, and lower half is ordered by their sequence similarities. Colors indicate biophysical group assignments and are assigned based on cophenetic distance. Clustering thresholds indicated on each dendrogram. Domains that cluster together based on sequence identity colored dark gray on dendrogram.

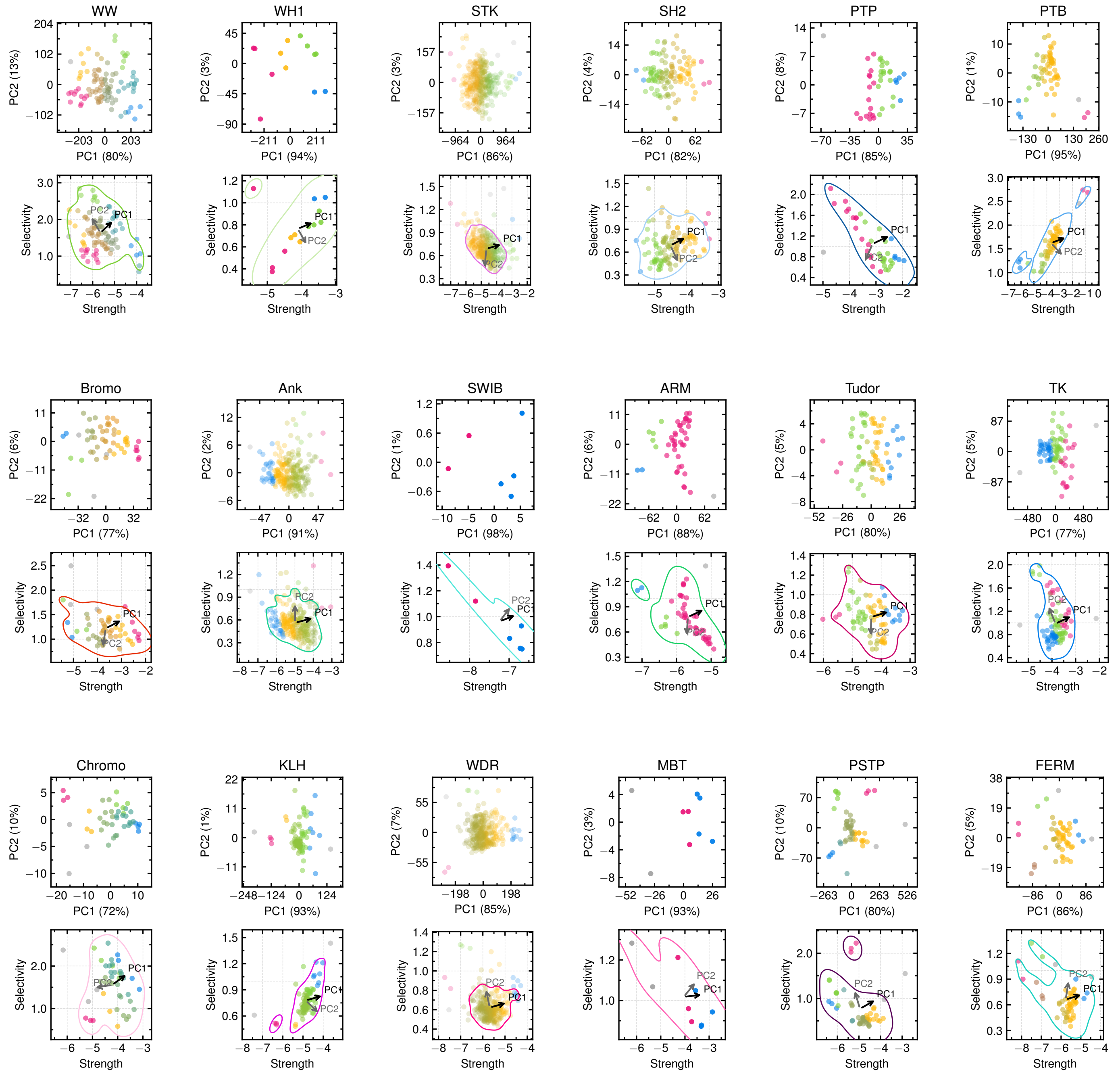


**Fig. S9 Biophysical equivalence groups capture quantitative differences in PBD binding strength and selectivity.** (Top plot) PCA of PBD–peptide affinities. PBDs projected onto first 2 PCs (percent variance given in parentheses) and colored by biophysical group. (Bottom plot) Binding strengths and selectivities of PBDs. Contours outline 90% of the probability mass per family. Arrows indicate directions of first two PCs determined through least squares regression.

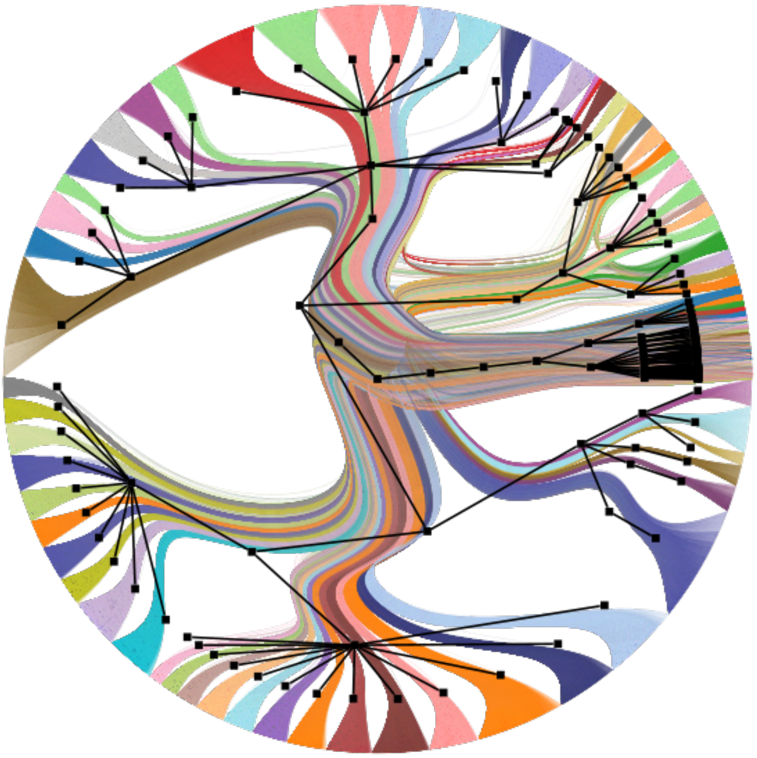

**Fig. S10 Graph representation of the SBM fit to QSM/N.** A full representation of the fitted SBM with proteins shown at the perimeter and colored by their block assignment in the finest resolution of the hierarchy. The internal graph shows the hierarchical structure of the SBM with connections between the lowest level blocks and individual proteins omitted.

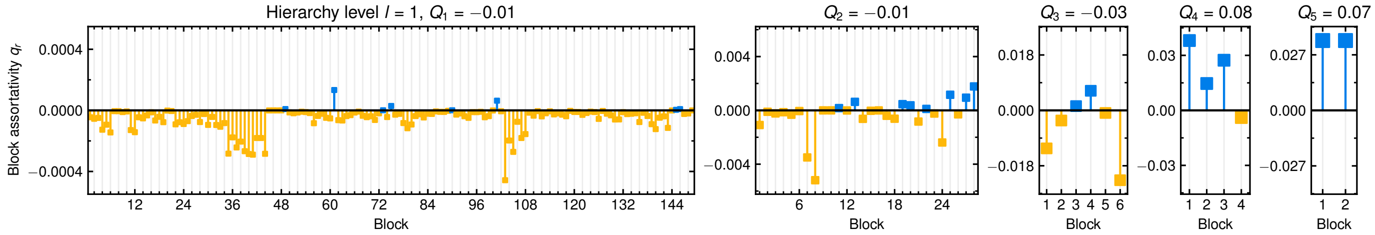


**Fig. S11 QSM/N contains both assortative and disassortative modules.** Local assortativity ($q_r$) of each block (module) of the 5-level hierarchical stochastic block model fit to QSM/N. Each level of the hierarchy is indexed by $l$, with the finest resolution structure of the network resolved in level 1, and block index is arbitrarily assigned. Newman modularity of each level ($Q_l$) is also reported. Local assortativity is computed as $q_r = \frac{1}{2E}\left(e_{rr} - \frac{e_r^2}{2E}\right)$, where $e_r r$ is twice the sum of edges within the block, $e_r = \sum_s e_{rs}$ is the sum of edges from block $r$ to blocks $s$, and $E$ is the sum of all edges.[75,76] With this definition of local assortativity, Newman modularity is computed as $Q_l = \sum_r q_r$.

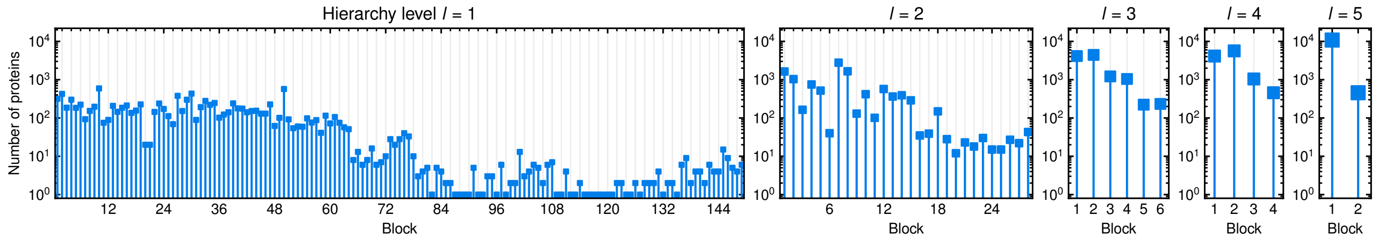


**Fig. S12 QSM/N contains modules of varying size.** Number of proteins assigned to each block (module) of the 5-level hierarchical stochastic block model fit to QSM/N. Each level of the hierarchy is indexed by $l$, with the finest resolution structure of the network resolved in level 1, and block index is arbitrary.

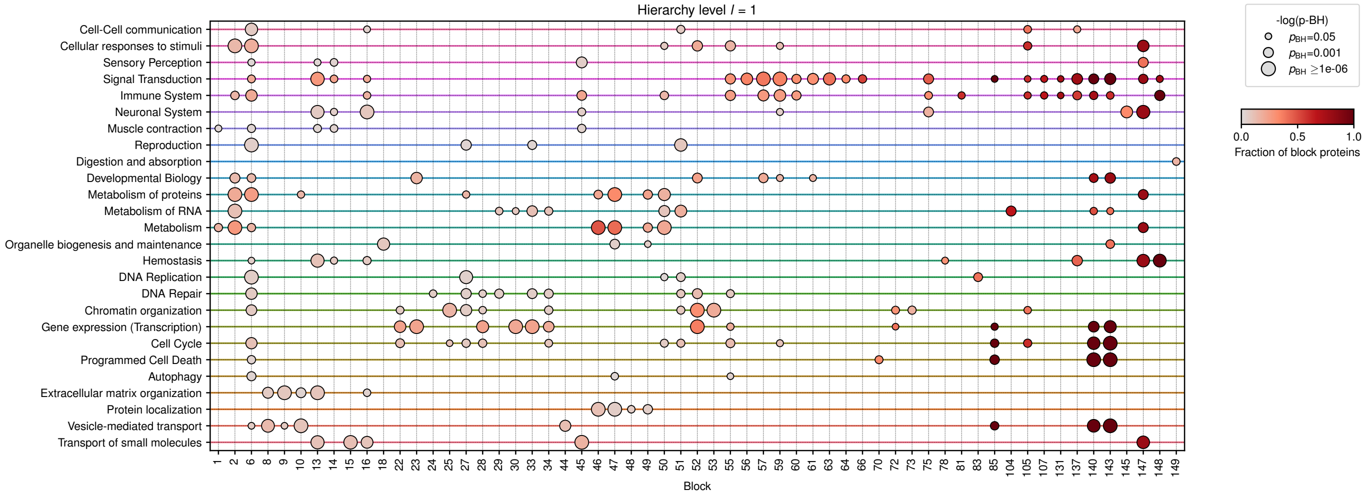


**Fig. S13 QSM/N contains modules enriched for known biological processes.** Dot plot indicates blocks (modules) in level 1 of the hierarchical SBM fit to QSM/N that show significant enrichment (5% Benjamini-Hochberg corrected FDR) for Reactome root pathways.[65] Dot size indicates significance of enrichment. Dot color indicates fraction of proteins in the block assigned to the enriched Reactome root pathway. Blocks without any enrichment are not shown.

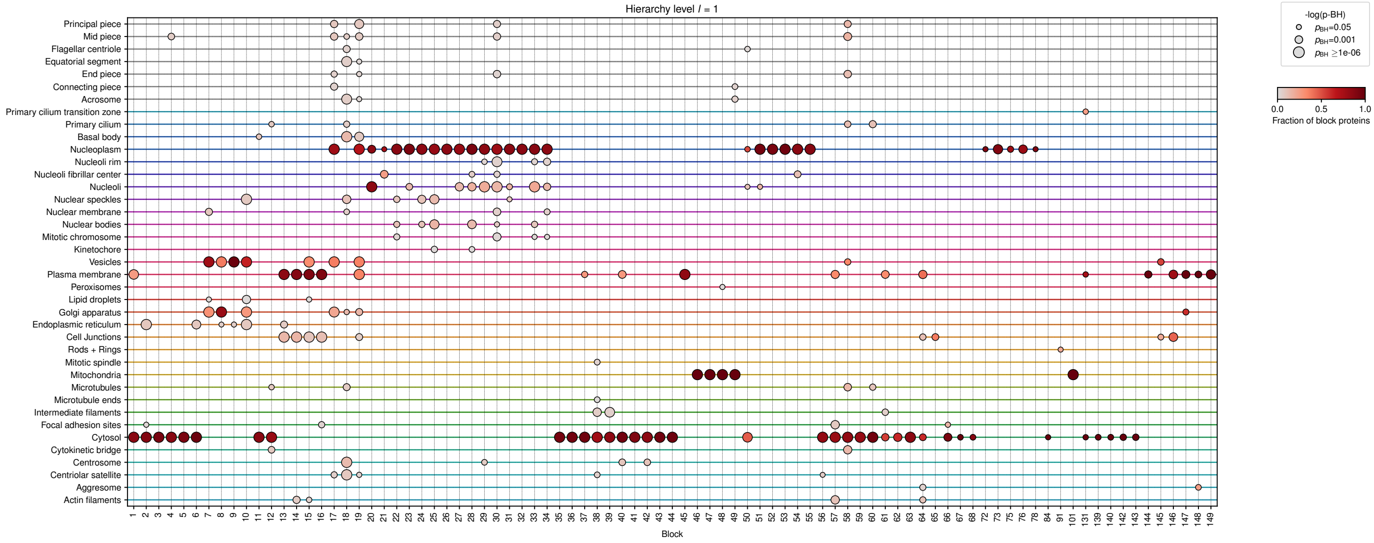


**Fig. S14 QSM/N contains modules enriched for specific subcellular localizations.** Dot plot indicates blocks (modules) in level 1 of the hierarchical SBM fit to QSM/N that show significant enrichment (5% Benjamini-Hochberg corrected FDR) for subcellular localization as determined in the Cell Atlas of the Human Protein Atlas[62]. Dot size indicates significance of enrichment. Dot color indicates fraction of proteins in the block assigned to the enriched subcellular localization. Blocks without any enrichment are not shown.

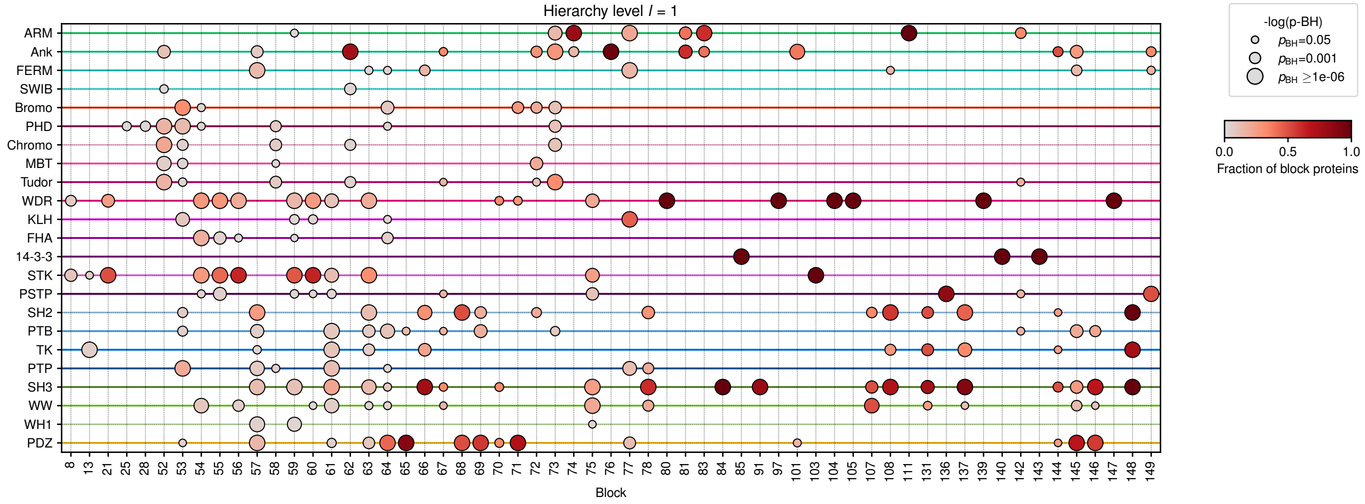


**Fig. S15 QSM/N contains modules enriched for specific PBD families.** Dot plot indicates blocks (modules) in level 1 of the hierarchical SBM fit to QSM/N that show significant enrichment (5% Benjamini-Hochberg corrected FDR) for a PBD family. Dot size indicates significance of enrichment. Dot color indicates fraction of proteins in the block assigned to the enriched PBD family. Blocks without any enrichment are not shown.

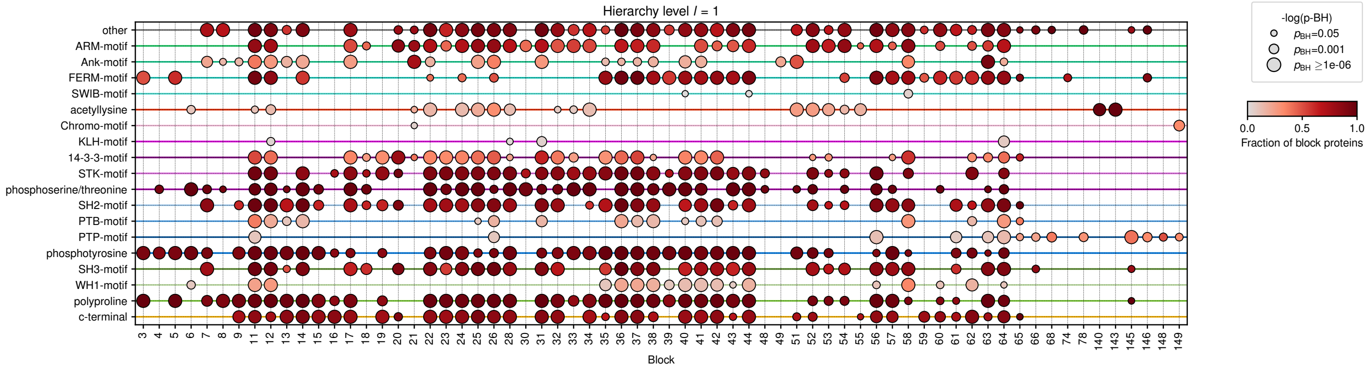


**Fig. S16 QSM/N contains modules enriched for specific peptidic binding chemistries.** Dot plot indicates blocks (modules) in level 1 of the hierarchical SBM fit to QSM/N that show significant enrichment (5% Benjamini-Hochberg corrected FDR) for a peptidic type. Dot size indicates significance of enrichment. Dot color indicates fraction of proteins in the block assigned to the enriched peptidic type. Blocks without any enrichment are not shown.

### Supplementary Methods

### Experimental measurement of PDZ–peptide affinities

#### Protein expression and purification

DNA encoding the PDZ domains from SHANK3 (Uniprot Q9BYB0, residues 636-741), DLG1-2 (Uniprot Q12959, residues 311-407), and DLG2-2 (Uniprot Q15700, residues 185-281) was obtained from Twist Bioscience and cloned into a kanamycin-resistant pET vector as a C-terminal fusion to $His_6$-SUMO using Gibson assembly. To express the recombinant fusion proteins, E. coli BL21(DE3) cells were transformed with the appropriate pET-$His_6$-SUMO-PDZ plasmid. Cells were grown at 37 °C in terrific broth supplemented with 50 $\mu$g/mL kanamycin until cells reached an optical density at 600 nm ($OD_{600}$) of 0.5. Then, isopropyl-$\beta$-D-1-thiogalactopyranoside (IPTG) was added to a final concentration of 0.5 mM to induce protein expression, and the cells were incubated at 18 °C overnight.

Cultures were centrifuged to pellet cells, which were then resuspended in lysis buffer (50 mM Tris pH 8.0, 300 mM NaCl, 10% glycerol, 10 mM imidazole, and freshly added 2 mM $\beta$-mercaptoethanol) with protease inhibitor cocktail [200 $\mu$M AEBSF (Calbiochem), 20 $\mu$M leupeptin (Calbiochem), 1 $\mu$M peptstatin A (Sigma)]. The cell suspensions were sonicated (Fisherbrand Probe Sonicator) to lyse cells. Lysates were centrifuged at 14,000xg for 45 min to pellet debris, then the supernatant was filtered through a 1.1 $\mu$m syringe filter. The filtrate was loaded onto a 5 mL HisTrap Ni-NTA column (Cytiva) and washed with 50 mL of lysis buffer (50 mM Tris pH 8.0, 300 mM NaCl, 10% glycerol, 10 mM imidazole, and freshly added 2 mM $\beta$-mercaptoethanol), followed by 50mL of wash buffer (50 mM Tris, pH 8.5, 50 mM NaCl, 10% glycerol, 10 mM imidazole, and freshly added 2 mM $\beta$-mercaptoethanol). The protein was eluted off the Ni-NTA column in elution buffer (50 mM Tris, pH 8.5, 50 mM NaCl, 500 mM imidazole, and 10% glycerol) and loaded onto a 5 mL HiTrap Q Anion exchange column (Cytiva). The column was washed using Anion A buffer (50 mM Tris, pH 8.5, 50 mM NaCl, 1 mM TCEP, and 10% glycerol), then proteins were eluted with a gradient between Anion A buffer and Anion B buffer (1 M NaCl, 50 mM Tris, pH 8.5, 1 mM TCEP, and 10% glycerol). Finally, the proteins were purified by size-exclusion chromatography on a Superdex 75 16/600 gel filtration column (Cytiva) equilibrated with SEC buffer (10 mM HEPES, pH 7.5, 150 mM NaCl, 1 mM TCEP, and 10% glycerol). Pure fractions were pooled and concentrated (3 kDa MWCO Millipore Amicon Ultra Centrifugal Filters). The protein was aliquoted in 200 $\mu$L aliquots and flash frozen in liquid N2 for long-term storage at -80 °C.

#### Fluorescence polarization experiments

FITC-labeled peptides were obtained from Synpeptide as lyophilized solids. Peptides were dissolved in 100 mM Tris, pH 8, and their concentrations were determined by measuring absorption at 495 nm and using the extinction coefficient for fluorescein. Working aliquots of each peptide stock were diluted down to 100 nM in buffer and stored at 4 °C, protected from light. Serial dilutions of each PDZ domain were prepared in FP Assay buffer (60 mM HEPES, pH 7.2, 75 mM KCl, 75 mM NaCl, 1 mM EDTA, and 0.05% Tween 20) and combined 1:1 v/v with each fluorescent peptide in a 384-well black plate. The final concentration of peptide was 50 nM and the final concentration of the PDZ protein ranged from the tens of nanomolar to the hundreds of millimolar (48.8 nM, 97.7 nM, 195.5 nM, 391 nM, 781 nM, 1.6 $\mu$M, 3.1 $\mu$M, 6.3 $\mu$M, 12.5 $\mu$M, 25 $\mu$M, 50 $\mu$M, 100 $\mu$M, 200 $\mu$M, 400 $\mu$M, and 800 $\mu$M). The plate was centrifuged at 4000g for 5 minutes and then incubated at room temperature, protected from light, for an hour to let the samples equilibrate. Parallel and perpendicular measurements were taken using the 485/530 polarization cube on the BioTek Neo2 Plate Reader. For each titration, a minimum of three separate replicates were carried out. Each replicate titration was individually fit to the quadratic binding equation:

$$A = r[PL] - r[L]$$

$$Y = r[L] + \frac{A\left([L]_t + K_\text{D} + X - \sqrt{-4[L]_t X + ([L]_t + K_\text{D} + X)^2}\right)}{2[L]_t},$$

where $[L]_t$ was constrained to 0.050 (50 nM, the concentration of the fluorescent peptide). The average and standard deviation of all replicates is reported.

#### Protein sequences

His-SUMO-DLG1-2:
MGSSHHHHHHGSGLVPRGSASMSDSEVNQEAKPEVKPEVKPETHINLKVSDGSSEIFFKIKKTTPLRRLMEAFAKRQGKE
MDSLRFLYDGIRIQADQTPEDLDMEDNDIIEAHREQIGGKPVSEKIMEIKLIKGPKGLGFSIAGGVGNQHIPGDNSIYVT
KIIEGGAAHKDGKLQIGDKLLAVNNVCLEEVTHEEAVTALKNTSDFVYLKVAKPTS

His-SUMO-DLG2-2:
MGSSHHHHHHGSGLVPRGSASMSDSEVNQEAKPEVKPEVKPETHINLKVSDGSSEIFFKIKKTTPLRRLMEAFAKRQGKE
MDSLRFLYDGIRIQADQTPEDLDMEDNDIIEAHREQIGGRPILETVVEIKLFKGPKGLGFSIAGGVGNQHIPGDNSIYVT
KIIDGGAAQKDGRLQVGDRLLMVNNYSLEEVTHEEAVAILKNTSEVVYLKVGKPTT

His-SUMO-SHANK3:
MGSSHHHHHHGSGLVPRGSASMSDSEVNQEAKPEVKPEVKPETHINLKVSDGSSEIFFKIKKTTPLRRLMEAFAKRQGKE
MDSLRFLYDGIRIQADQTPEDLDMEDNDIIEAHREQIGGSHSDYVIDDKVAVLQKRDHEGFGFVLRGAKAETPIEEFTPT
PAFPALQYLESVDVEGVAWRAGLRTGDFLIEVNGVNVVKVGHKQVVALIRQGGNRLVMKVVSVTR

**Peptide sequences**

PDZ 1: FITC-PEG2-AHFSSLESEV-COOH
PDZ 2: FITC-PEG2-PQAQAENTAF-COOH
PDZ 3: FITC-PEG2-QIAGTKSTTV-COOH
PDZ 4: FITC-PEG2-TTVRCLEAEV-COOH
PDZ 5: FITC-PEG2-STLTIFETAL-COOH
PDZ 6: FITC-PEG2-SHSSKGETAV-COOH
PDZ 7: FITC-PEG2-EPPEEFDTAL-COOH
PDZ 8: FITC-PEG2-NQLAWFDTDL-COOH
PDZ 9: FITC-PEG2-DFREDDDTAL-COOH
PDZ 10: FITC-PEG2-SGSLKVMTTV-COOH
PDZ 11: FITC-PEG2-TRVVDQITTV-COOH
PDZ 12: FITC-PEG2-LRKPEADTAL-COOH
